# PIKfyve influences inter-organelle contacts with lysosomes to modulate the endoplasmic reticulum

**DOI:** 10.1101/2025.07.15.664974

**Authors:** Nicala Jenkins, Zainab Adamji, Maria R. Narciso, Nourhan R. Almasri, Sharifa Lomiyev, Roberto J. Botelho

**Author notes:** To whom correspondence should be sent.

## Abstract

Lysosomes clear unwanted cellular material delivered by constant membrane fusion. Membrane fission is thus required to balance lysosome size, number, and composition. PIKfyve is a lipid kinase that converts phosphatidylinositol-3-phosphate [PtdIns(3)P] to phosphatidylinositol-3,5-bisphosphate [PtdIns(3,5)P_2_] and promotes lysosome fission since lysosomes coalesce into larger, but fewer organelles in its absence. Here, we reveal a role for PIKfyve in regulating endoplasmic reticulum (ER) dynamics. We show the ER is less reticulated and motile in cells inhibited for PIKfyve. Partly, this arises because lysosomes cluster perinuclearly and are less motile, which appears to arrest ER hitchhiking, a process in which lysosomes pull and form ER tubules. Secondly, the ER morphology is distorted because of hyper-tethering of protrudin, an ER transmembrane protein, to lysosomes via excess PtdIns(3)P and protrudin’s FYVE domain. Our findings reveal that PIKfyve balances phosphoinositides at ER-lysosome contact sites to govern ER properties and have significant implications for our understanding of PIKfyve function and of diseases linked to its dysfunction.

**Summary:** Long associated with the regulation of the endolysosomal pathway, the authors show that the PIKfyve lipid kinase extends its sphere of influence to modulate the architecture and dynamics of the endoplasmic reticulum by balancing phosphoinositide species at their inter-organellar sites.

## Introduction

Lysosomes are acidic and degradative organelles that receive and digest a plethora of molecular cargo from the endocytic, autophagic and phagocytic pathways (Ballabio and Bonifacino, 2020; Inpanathan and Botelho, 2019). They are also highly dynamic because they continuously undergo fusion-fission cycles with maturing late endosomes, transport vesicles from the *trans*-Golgi network, and between themselves (Saffi and Botelho, 2019; Luzio et al., 2007). This constant intermixing is dependent on bidirectional movement of lysosomes on microtubules driven by kinesin and dynein motors (Cabukusta and Neefjes, 2018). Perhaps due to this dynamic nature, lysosomes are a heterogeneous collection of organelles that differ in shape, size, position, and function (Saffi and Botelho, 2019; Inpanathan and Botelho, 2019). Thus, and unless otherwise indicated, we use the term *lysosomes* to encapsulate this full spectrum, including late endosomes, terminal lysosomes, and their hybrids, endolysosomes.

Given this complexity, lysosomes are subject to various regulatory inputs including by the Rab7 and Arl8b GTPases and the PIKfyve lipid kinase (Ballabio and Bonifacino, 2020; Bissig et al., 2017; de Lartigue et al., 2009; Cantalupo et al., 2001; Khatter et al., 2015). PIKfyve is a conserved lipid kinase that localizes to endosomes and lysosomes via its FYVE domain and phosphatidylinositol-3-phosphate [PtdIns(3)P], which is then converted by PIKfyve into phosphatidylinositol-3,5-bisphosphate [PtdIns(3,5)P_2_] (Ikonomov et al., 2001; Sbrissa et al., 2002, 2007). While deletion of PIKfyve is embryonic lethal, tissue-specific knockouts or mutations that abate PtdIns(3,5)P_2_ levels are associated with defects in immunity, inflammation, metabolic, and neurological functions (Ikonomov et al., 2011; Min et al., 2014; Chow et al., 2007; Tsuruta et al., 2009; Cheng et al., 2025; Min et al., 2019; Prado et al., 2025). At the molecular level, there remain few known PtdIns(3,5)P_2_-binding proteins, which include Ca^2+^, Cl^-^ and Na^+^ channels like TRPML1, ClC7, and TPCs, respectively (Leray et al., 2022; Dong et al., 2010; Wang et al., 2012), the actin regulator cortactin (Hong et al., 2015), and membrane curvature factors like the yeast Atg18 (Gopaldass et al., 2017). Through these and other unknown effectors, PIKfyve and PtdIns(3,5)P_2_ collectively modulate osmosis, ion-based signaling, lysosome fusion-fission cycling, and the degradative capacity of lysosomes (Bissig et al., 2017; Choy et al., 2018; Leray et al., 2022; Dong et al., 2010; Sharma et al., 2019; Wang et al., 2012). Arguably, the most conspicuous defect caused by PIKfyve loss of function is lysosome enlargement and the concomitant decline in lysosome number, likely due to reduced membrane fission relative to fusion (Bissig et al., 2017; Choy et al., 2018; Sharma et al., 2019). However, we do not know if this occurs by dysregulation of fission complexes and/or of signaling factors like ion-based signaling. Interestingly, endosomes and mitochondrial sites of fission are demarcated by endoplasmic reticulum (ER)-dependent contacts (Friedman et al., 2013; Abrisch et al., 2020; Friedman et al., 2011). We thus considered whether PIKfyve and the ER may play a joint role in lysosome fission.

The ER is a reticulated membrane network composed of membrane sheets and tubules extending throughout the cell, and forming functional domains such as the rough ER and smooth ER (Westrate et al., 2015). The ER morphology is regulated by various proteins and responds to signals such as the unfolded protein response and the nutrient status of the cell (Chen et al., 2023; Goyal and Blackstone, 2013; Lu et al., 2020). For instance, ER sheets are promoted by integral membrane proteins such as CLIMP63, which form bridge-like structures connecting opposing ER bilayers (Shibata et al., 2010). In comparison, ER tubules are dynamic, continually forming, branching, and rearranging through the action of proteins such as Reticulons and Atlastins (Wang et al., 2016; Hu et al., 2009; Voeltz et al., 2006; Goyal and Blackstone, 2013).

ER remodeling can also occur through motor and cytoskeleton-driven processes. This includes microtubule tip attachment complex (TAC)-mediated movement and ER-sliding, which rely on the interaction between the ER membrane and microtubules to facilitate the formation of new ER tubules (Waterman-Storer and Salmon, 1998; Rodríguez-García et al., 2020; Spits et al., 2021), and ER hitchhiking. During ER hitchhiking, the ER forms a contact with vesicles, endosomes, or lysosomes (Guimaraes et al., 2015; Spits et al., 2021; Raiborg et al., 2015b; Lu et al., 2020; Langley et al., 2025). The attached ER then co-migrates with the motile endosome/lysosome in a motor and microtubule-dependent manner to grow a new ER tubule (Guimaraes et al., 2015; Spits et al., 2021; Voeltz et al., 2024; Lu et al., 2020). Little is known about the protein complexes holding the ER-endosome/lysosome together, but these are thought to be maintained by tethering proteins such as protrudin (Raiborg et al., 2015b; a; Pedersen et al., 2020; Spits et al., 2021).

Protrudin, which was originally identified as a protein that promotes neurite formation (Shirane and Nakayama, 2006), is an ER-transmembrane protein that together with VAPA, another ER transmembrane protein, and the lysosomal Rab7 GTPase, form an ER-lysosome tethering complex (Raiborg et al., 2015a; Pedersen et al., 2020; Matsuzaki et al., 2011; Saita et al., 2009). Protrudin has a FYVE domain and a protein-binding low complexing region (LCR) that respectively are thought to bind to PtdIns(3)P and GTP-bound Rab7 on endosomes/lysosomes (Raiborg et al., 2015a; Pedersen et al., 2020). Protrudin then extends its cytoplasmic region to loop around and interact with VAPA via its FFAT motif. Through this contact, protrudin is thought to load kinesin-1 onto lysosome-bound FYCO1, a Rab7 effector that promotes lysosome anterograde movement and binds PtdIns(3)P (Pedersen et al., 2020; Raiborg et al., 2015a; Jongsma et al., 2024; Matsuzaki et al., 2011; Pankiv et al., 2010). Collectively, protrudin tethers ER-lysosomes together and modulates their relative dynamics (Cabukusta and Neefjes, 2018; Jongsma et al., 2016, 2024).

Overall, there is a bidirectional relationship between the ER and organelles of the endo-lysosomal pathway through contact sites. ER contacts appear to define areas of endosome fission, while endosomes and lysosomes help reshape the ER via hitchhiking. Given this, we postulated that ER-driven fission may malfunction during PIKfyve inhibition, resulting in lysosome coalescence. Additionally, we hypothesized that lysosome coalescence during PIKfyve loss may alter ER morphology and dynamics. Here, we provide evidence that PIKfyve inhibition potently distorts the ER morphology and dynamics by hindering ER hitchhiking and stabilizing protrudin interaction with lysosomes via excess PtdIns(3)P levels.

## Results

### Lysosome morphology is relatively intact in cells manipulated for ER morphology

PIKfyve inhibition enlarges lysosomes via coalescence, ostensibly by perturbing fission (Choy et al., 2018; Bissig et al., 2017; Sharma et al., 2019). On the other hand, sites of tubular endosome fission are demarcated by ER tubules that wrap around endosomes (Friedman et al., 2013; Rowland et al., 2014). Conceivably then, lysosome coalescence in PIKfyve-inhibited cells may be the corollary of perturbed contacts between ER and endo-lysosomal structures. To test this hypothesis, we over-expressed CLIMP63 or Rtn4A, which respectively bias ER towards sheets and reticulated tubules (Shibata et al., 2010; Wang et al., 2016; Wu and Voeltz, 2021; Voeltz et al., 2006), and examined their effect on the morphology and number of lysosomes labelled with fluorescent dextrans in resting and apilimod-treated cells (Choy et al., 2018; Ohkuma and Poole, 1978; Ferris et al., 1987).

As expected, COS-7 cells overexpressing mCh-CLIMP63 displayed an overwhelmingly sheet-like ER network with a dramatic decline of ER tubules **(Fig. 1A).** Conversely, cells expressing eGFP-Rtn4 had an exaggerated tubular ER network **(Fig 1B).** We then quantified the total lysosome number and the individual lysosome volume in cells over-expressing CLIMP63 or Rtn4 and treated with vehicle or apilimod to block PIKfyve. Remarkably, there was no difference in lysosome size and number in cells expressing CLIMP63 or Rtn4 compared to the respective mCherry or GFP controls **(Fig. 1C-F).** This shows that distortion of ER morphology does not cause lysosome coalescence and does not mimic PIKfyve inactivation. Interestingly, overexpression of CLIMP63 or Rtn4 respectively enhanced and reduced apilimod-driven changes in the volume of individual lysosomes **(Fig. 1D, 1F),** but did not alter the total lysosome number per cell (**Fig. 1C, 1E**). This suggests that the ER morphology can impact lysosome size, but not by coalescence as in PIKfyve inactivation. Overall, these results show that manipulating ER morphology does not readily mimic the effects of PIKfyve on lysosome dynamics but can synergize with PIKfyve inactivation to influence lysosome size.

**Figure 1:**
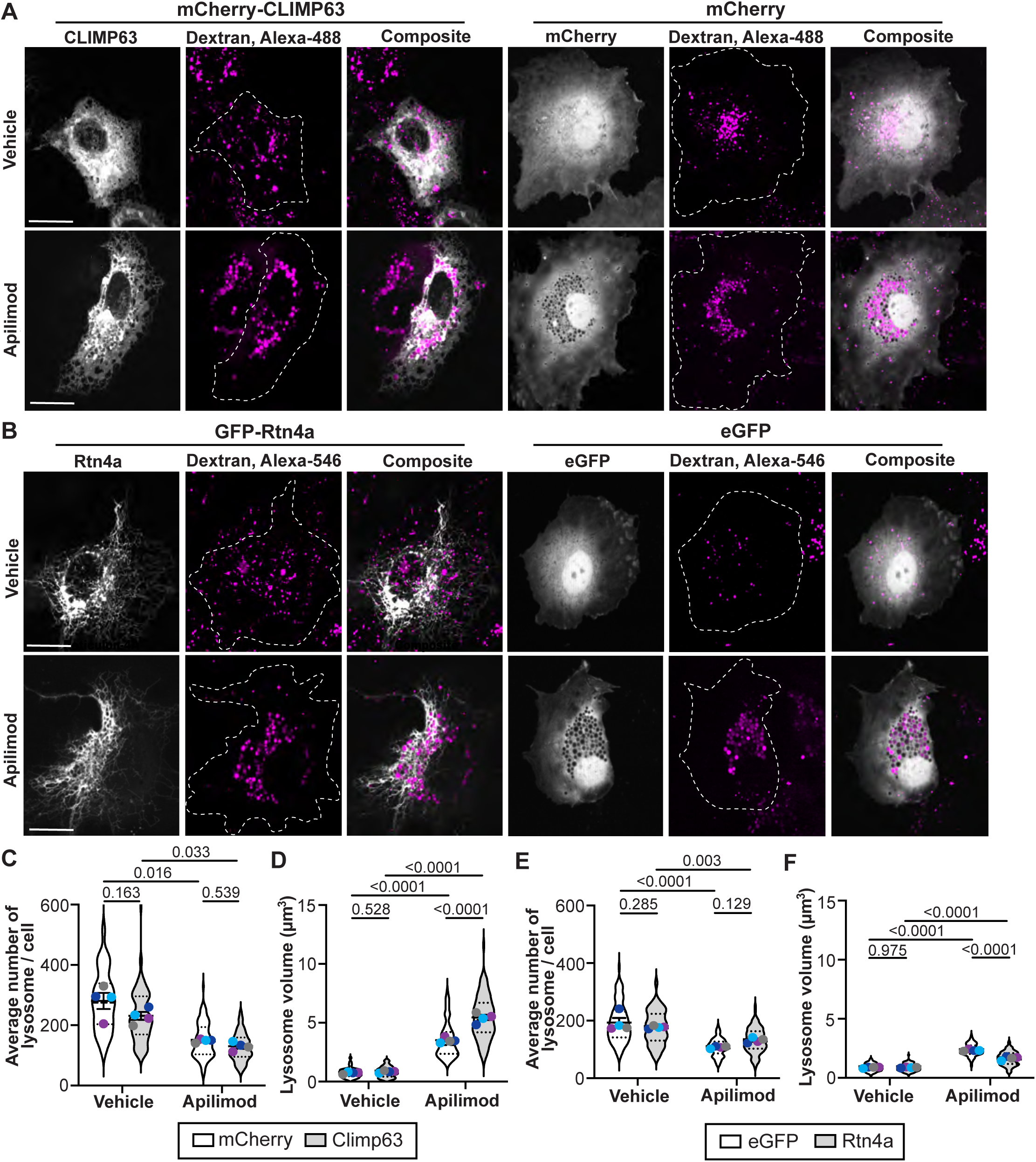
Overexpression of ER morphogenic proteins does not mimic PIKfyve inhibition in COS-7 cells. **(A, B)** Confocal images of COS-7 cells labeled with fluorescent dextran (magenta) and transiently expressing mCherry-CLIMP63 or mCherry (gray; A), or eGFP-Rtn4a or eGFP (gray; B). Cells were exposed to either 0.01% DMSO (vehicle) or 80 nM apilimod for 2 h. Scale bar = 20 µm. **(C-D)** Quantification of lysosome number (C) and volume (D) upon expression of CLIMP63 to induce ER sheets. **(E-F)** Quantification of lysosome number (E) and volume (F) upon expression of Rtn4a to promote ER tubulation. All experiments were repeated four independent times. Data points from matching independent experiments are colour coded. Shown are the mean ± SEM. Data are based on 30-50 transfected cells per condition per experiment and analyzed using repeated measures two-way ANOVA and Tukey’s multiple comparisons test; p values are shown.

### PIKfyve inhibition distorts ER morphology

We next considered the converse: whether lysosome distortion in PIKfyve-inhibited cells would alter the ER morphology. To test this, we expressed ER-mNeonGreen in COS-7 cells to label the ER and labelled lysosomes with a fluorescent dextran. We opted to first use a pharmacological approach to block PIKfyve using apilimod or YM-201636 because this permitted acute disturbance of lysosomes, mitigating indirect effects on the ER.

In vehicle-treated cells, the ER network exhibited a series of sheets and tubules throughout the cell cytoplasm, some of which were juxtaposed to dextran-labelled lysosomes **(Fig. 2A, Sup. Fig. S1A).** Strikingly, the ER network was visibly distorted, exhibiting numerous “black holes” in cells treated with apilimod **(Fig. 2A)** and YM-201636 **(Sup. Fig. S1A).** Many of these voids, but not all, were occupied with Alexa647-dextran **(Fig. 2A, Sup. Fig. S1A).** To determine if some of these voids were cytosolic in nature or contoured by other swollen early endosomal structures, we expressed cytosolic mCherry or mCherry-Rab5. None of the apparent ER-decorated voids labelled with cytosolic mCherry (**Sup. Fig. 2A)**, but a subpopulation was lined with mCherry-Rab5 **(Sup. Fig. S2B).** Overall, the ER “black holes” were occupied with either dextran-labelled lysosomes or mCherry-Rab5 endosomes **(Sup. Fig. S2C-D)**, indicating that separate populations of enlarged endosomal and lysosomal organelles may alter the ER architecture. To complement our visual inspection and buttress these assertions, we quantified the ER morphology by extracting total ER skeletal length, number of branches and junctions per cell, and average branch length (depicted in **Fig. 2B, 2C**). Apilimod and YM201636 both caused a significant decline in ER total skeletal length, loss of ER junctions and branch numbers, though not for ER branch length **(Fig. 2D-G** and **Sup. Fig. S1B-E).** Importantly, these changes were not due to an acute and transient effect of PIKfyve inhibition since cells silenced for *VAC14* for 48 h displayed the expected enlarged lysosomes and distorted ER morphometrics (**Sup. Fig. S3**). Collectively, these data show a reduction in total ER network size and loss of the ER interconnectivity in cells negated for PIKfyve activity.

**Figure 2:**
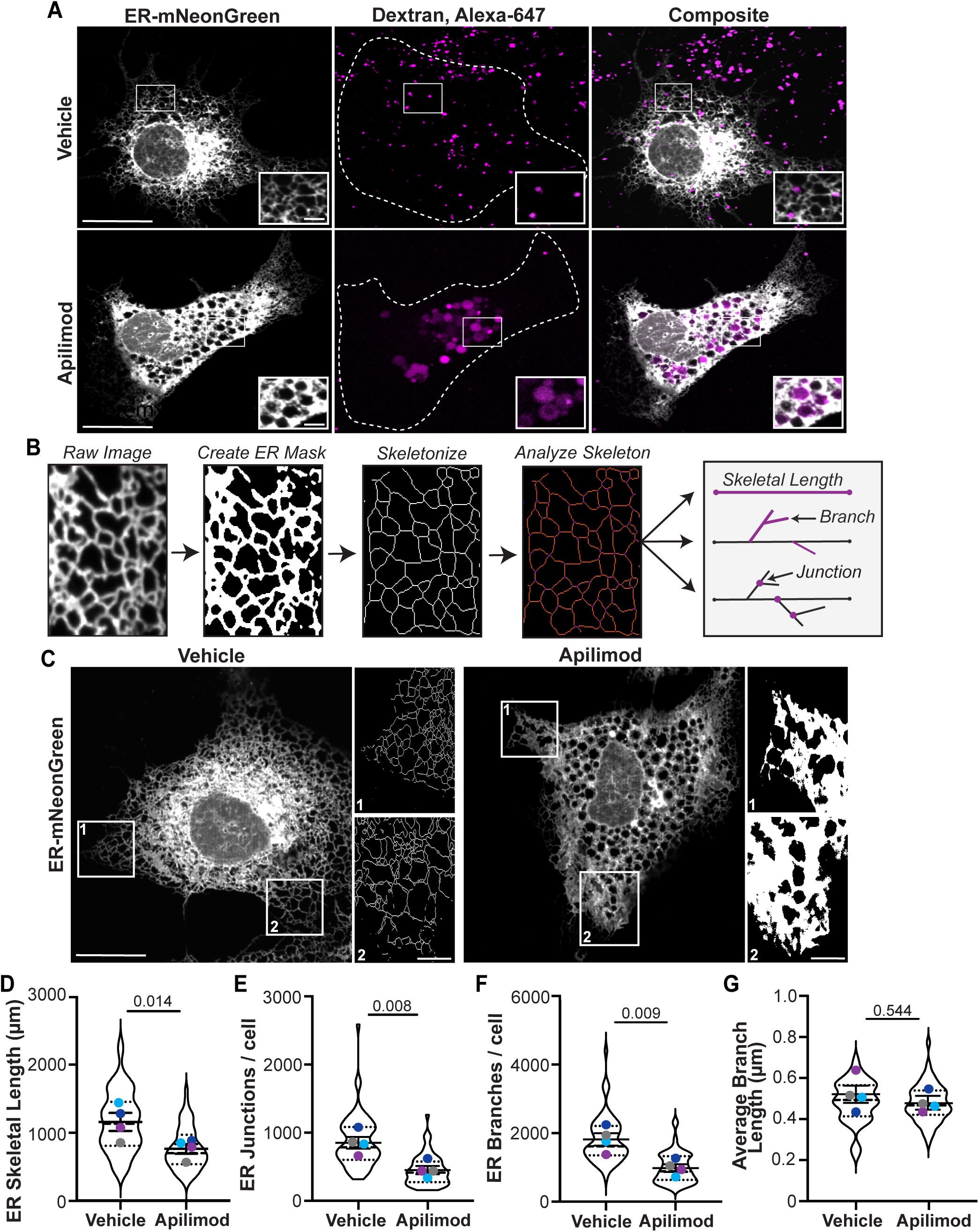
PIKfyve loss of function alters the ER architecture. **(A)** Confocal images of COS-7 cells transiently transfected with ER-mNeonGreen (gray), labeled with fluorescent Dextran (magenta), and treated with 0.01% DMSO (vehicle) or 80 apilimod for 2 h. **(B)** Image analysis strategy to quantify ER morphometrics whereby a fluorescence image is converted to an ER mask, skeletonized, and analysed for skeletal length, branch number, and junction number. **(C)** ER-mNeonGreen and corresponding skeletonized images of a cell treated with vehicle or apilimod. Scale bar: full size = 20 µm, zoom insert = 5 µm. **(D-G)** Quantification of ER morphology including ER skeletal length (D), number of ER junctions per cell (E), number of branches per cell (F), and average branch length (µm) (G). All experiments were repeated four independent times. Data points from matching independent experiments are colour coded. Shown are the mean ± SEM. Data are based on 15-20 transfected cells per condition per experiment. Data was analysed using a two-tailed paired Student’s t-test and p values are shown.

### Loss of PIKfyve activity impairs ER dynamics

We next examined whether ER motility was disturbed in PIKfyve-inhibited cells. To assess this, we employed a previously described method that measures membrane displacement by tracking pixel intensity changes over time (Spits et al., 2021). To achieve this, we tracked ER-mNeonGreen pixels over time in control and PIKfyve-hindered cells, binning pixel displacement into three categories: low (displacement of 0-3 pixels), intermediate (displacement of 4-7 pixels), and rapid ER movement (8 or more pixels; **Fig. 3A**). Relative to vehicle-treated cells, cells exposed to apilimod or YM201636 displayed a greater number of pixels with low membrane displacement and a corresponding drop in pixels with rapid membrane displacement **(Fig. 3B-E**; **Sup. Fig. S1F-I**; **Sup. Videos 1-3**). These results reveal that the ER is less motile in cells arrested for PIKfyve activity, further reinforcing the idea that lysosomal coalescence distorts ER architecture and dynamics.

**Figure 3:**
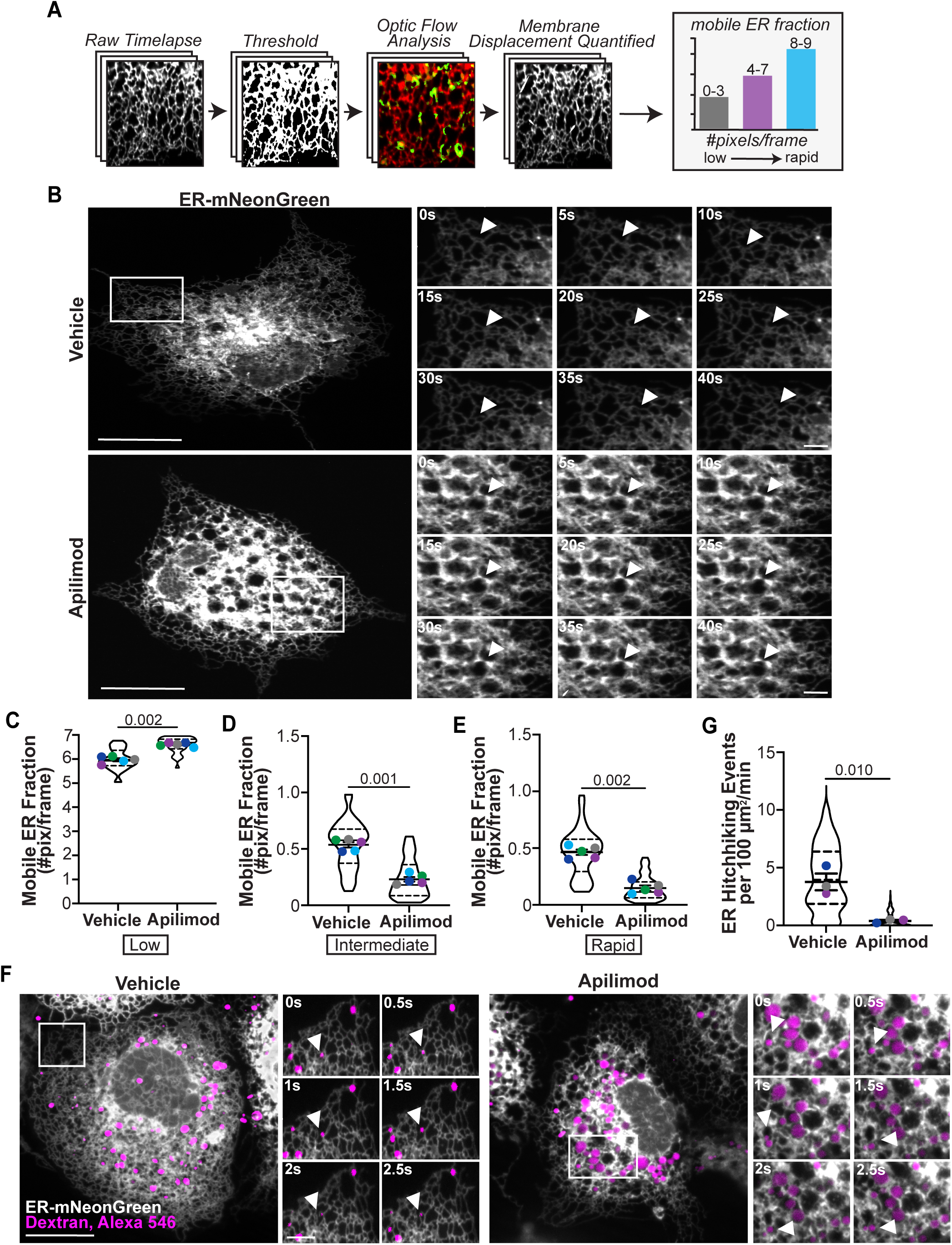
ER motility is reduced in PIKfyve inhibited cells. **(A)**Strategy to quantify ER motility from timelapse images, optic flow analysis and membrane displacement by grouping pixel changes per frame as indicated. **(B)** Representative time-lapse confocal images of COS-7 cells transiently transfected with ER-mNeonGreen (gray). Zoom inserts show ER redistribution as indicated by the white arrow. Cells were exposed to either 0.02% DMSO (vehicle) or 240 nM apilimod for 2 h. **(C-E)** Quantification of membrane displacement analysis categorized by low movement (C), intermediate movement (D), and rapid movement (E). **(F)** Confocal images of COS-7 cells transiently transfected with ER-mNeonGreen (gray), labeled with fluorescent dextran (magenta), and exposed to either 0.02% DMSO (vehicle) or 240 nM apilimod. Zoom inserts show an ER-hitchhiking event occurring over 2.5 seconds as indicated by the white arrow. Scale bar: full size = 20 µm, zoom insert = 5 µm. **(G)** Quantification of the number of ER-hitchhiking events that occurred in a 100 µm^2^ ROI over one minute, repeated three independent times. Data from *C-E* are from five independent experiments and based on 7 randomized ROIs from 7-10 transfected cells, per condition per experiment. Data in G are from three experiments and based on 5 randomized ROIs from 15-20 transfected cells, per condition per experiment. Data points from matched experiments are colour-coded. Shown is the mean ± SEM, analysed using a two-tailed paired Student’s t-test; p values are shown.

### ER hitchhiking is blunted in PIKfyve-perturbed cells

Conceivably, the distortion of the ER spatio-temporal properties in PIKfyve-deficient cells could simply arise from physical hindrance of swollen endo-lysosomes compressing the intricate ER network of tubules and sheets. While physical hindrance of swollen lysosomes may very well help disfigure the ER network, we noted that there were few swollen lysosomes in the periphery of PIKfyve-inhibited cells (quantified below), which nevertheless sustained loss of ER complexity. We thus postulated that a more sophisticated mechanism was at play.

The ER is partly sculpted by forming and extending new ER tubules, which can then fuse with existing tubules to promote ER interconnectivity (Westrate et al., 2015; Guo et al., 2018; Shibata et al., 2010; Spits et al., 2021). There are several processes that drive ER shaping that depend on microtubules: ER sliding, TAC-dependent ER extension, and ER hitchhiking (Spits et al., 2021; Grigoriev et al., 2008; Rodríguez-García et al., 2020; Guo et al., 2018; Shibata et al., 2010). We thus investigated whether perturbation of some of these processes contributed to changes in ER spatial-temporal dynamics in PIKfyve-negated cells.

First, using immunofluorescence against α-tubulin to analyze microtubule morphology, we did not perceive a significant difference in microtubule organization between vehicle and apilimod-treated cells **(Sup. Fig. S4A)** when quantifying parameters such as number of microtubule branches and junctions per cell, and average microtubule branch length **(Sup. Fig. S4B-D)** – this despite visibly enlarged lysosomes in apilimod-treated cells. This was also true for the number and kinetics of EB1-GFP comets, which associate with microtubule plus-ends **(Sup. Fig. S4E-H; Sup. Videos 4-6).** We then measured ER hitchhiking events by tracking ER and lysosome movement over one-minute movies at 0.3 Hz. Timelapses were then analysed for lysosome-ER co-migration in control and PIKfyve-inhibited cells. We observed an average of 4-6 hitchhiking events/min/100 µm^2^ area in control cells, which substantially dropped to 0-1 events/min/100 µm^2^ in apilimod-exposed cells **(Fig. 3F, G; Sup. Videos 7-8).** These observations support a model whereby the botched ER morphology and dynamics in PIKfyve-defective cells are partly underpinned by failed ER hitchhiking.

### Lysosome distribution and movement are altered in PIKfyve-inhibited cells

Hitchhiking requires the ER to attach to a endosome/lysosome and for the endosome/lysosome to move (Spits et al., 2021; Pedersen et al., 2020; Voeltz et al., 2024). We thus reasoned that abrogation of ER hitchhiking in PIKfyve-hindered cells may ensue from a defect in attachment and/or lysosome motility. We first tested lysosome motility and distribution in control and PIKfyve-impaired cells. Using shell analysis, we reveal that lysosomes collapse into the perinuclear region and become less frequent in the cell periphery in the absence of PIKfyve activity **(Fig. 4A-C).** This was accompanied by a significant decline in lysosome speed and total distance travelled in apilimod-treated COS-7 cells relative to vehicle-treated cells (**Fig. 4D-F),** made visible through kymographs that revealed a marked decrease in the motility of lysosomes upon PIKfyve inhibition (**Fig. 4G)**. Collectively, loss of PIKfyve activity is associated with hindered lysosome motility and displacement to the perinuclear region. This implies that in the absence of PIKfyve activity, lysosomes likely sustain a loss of kinesin and/or hyperactivation of dynein activities. As a corollary, this could impair ER hitchhiking and disrupt the ER network.

**Figure 4.**
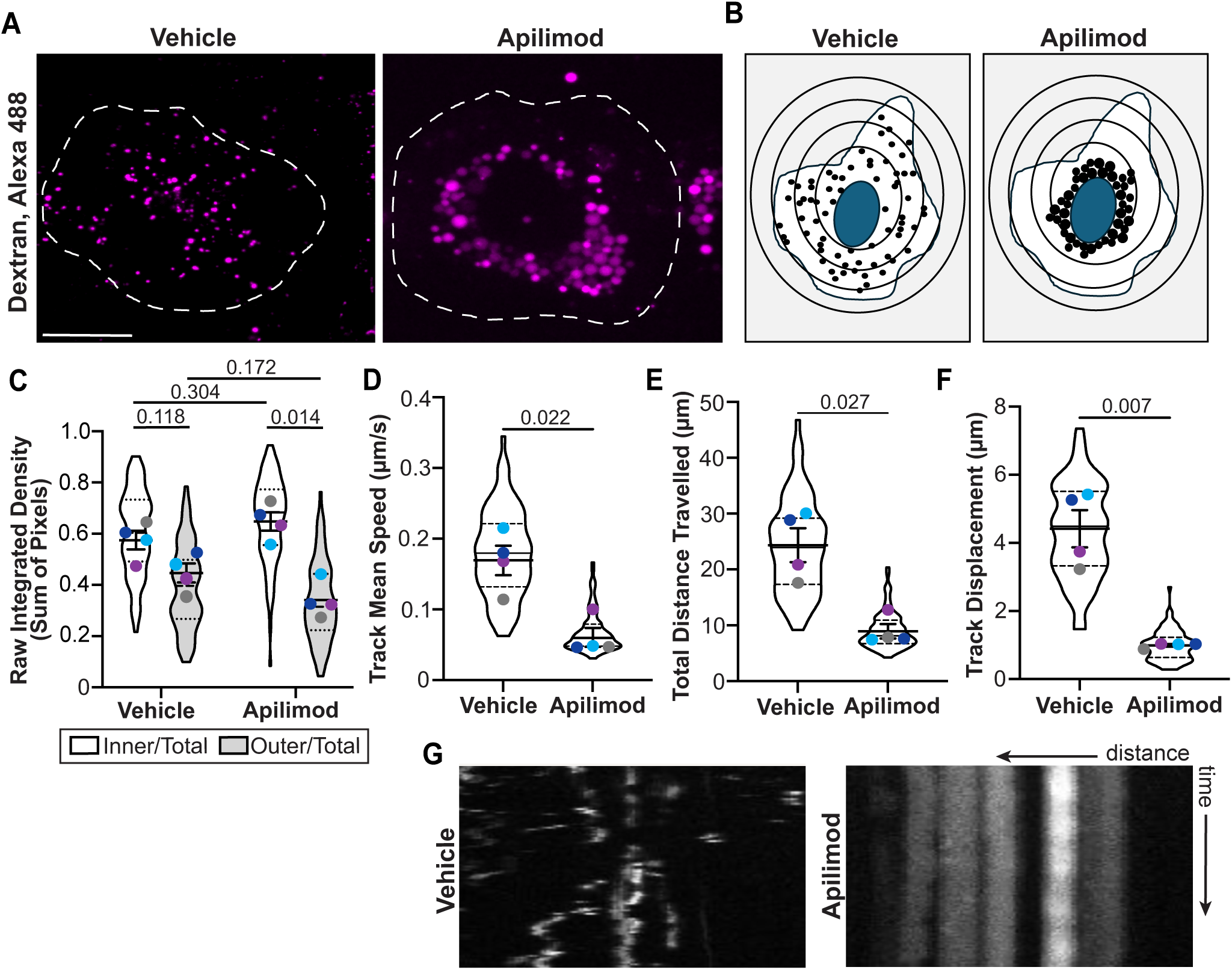
PIKfyve activity supports lysosome motility and spatial distribution. **(A)** Confocal images of COS-7 cells labeled with Alexa Fluor 647-Dextran (magenta) and exposed to either 0.02% DMSO (vehicle) or 240 nM apilimod for 2 h. Scale bar = 20 µm. **(B)** Schematic representation of the changes to lysosome spatial distribution between control and PIKfyve inhibited conditions. **(C)** Quantification of the distribution of lysosomes in the perinuclear region versus the peripheral region of the cell. A ratio of the number of lysosomes in the perinuclear region to total number of lysosomes in the cell was determined – the same was done for the peripheral region respectively. **(D-F)** Quantification of lysosome motility including track mean speed (D), total distance travelled (E), and track displacement (F). (**G)** Kymograph of lysosome movement along a linear ROI. The vertical axis represents time (s), and the horizontal axis represents distance along a selected path. Diagonal or segmented lines indicate motility; vertical lines represent stationary organelles. All experiments were repeated four independent times. Data points from matching independent experiments are colour coded. Shown is the mean ± SEM. Data were analysed using a two-way ANOVA and Tukey’s multiple comparison test for (C) and two-tailed paired Student’s t-test for (D-F); p values are shown.

### PIKfyve inhibition alters ER-lysosome contact sites

There are likely several types of ER-lysosome contact sites (Rudnik et al., 2024; Raiborg et al., 2015a; Allison et al., 2017; Du et al., 2023; Bonet-Ponce and Cookson, 2022; Cai et al., 2022; Gao et al., 2022; Saric et al., 2021). Nonetheless, contact sites formed by the ER transmembrane proteins, protrudin and VAPA/B, and the lysosomal Rab7 GTPase, are linked to lysosome and ER position and motility via kinesin (Pedersen et al., 2020; Raiborg et al., 2015a; Matsuzaki et al., 2011). Therefore, we tested if PIKfyve blockade altered the protrudin-VAPA-Rab7 contact site by using proximity ligation assays (PLAs) – representative figures of negative controls for each PLA pair are shown in **Sup. Fig. S5A-H**.

We found that protrudin and VAPA appeared proximal in equal measures between PIKfyve-enabled and PIKfyve-blocked cells **(Fig. 5A, D)**, which is perhaps not surprising since both are ER transmembrane proteins. However, we observed an inverse relationship between Rab7 and these proteins: there were fewer proximal events between VAPA and Rab7 (**Fig. 5B, E)**, but higher between protrudin and Rab7 **(Fig. 5C, F)**. Perhaps consistent with this, we did not observe a change in total Rab7 associated with LAMP1 by immunofluorescence in these cells and experimental conditions (**Sup Fig. S5I-K)**. Similarly, we did not observe a change in the cellular levels of GTP-loaded Rab7 as reported by affinity precipitation of GTP-bound Rab7 using GST-RILP-C33 in HeLa cells **(Sup Fig. S5L-M)**, though it is possible that specific pools of GTP-Rab7 may be altered in these conditions. This suggests that PIKfyve inactivation does not lead to a wholesale loss or enhancement of this membrane contact type but instead may be altering its conformation.

**Figure 5.**
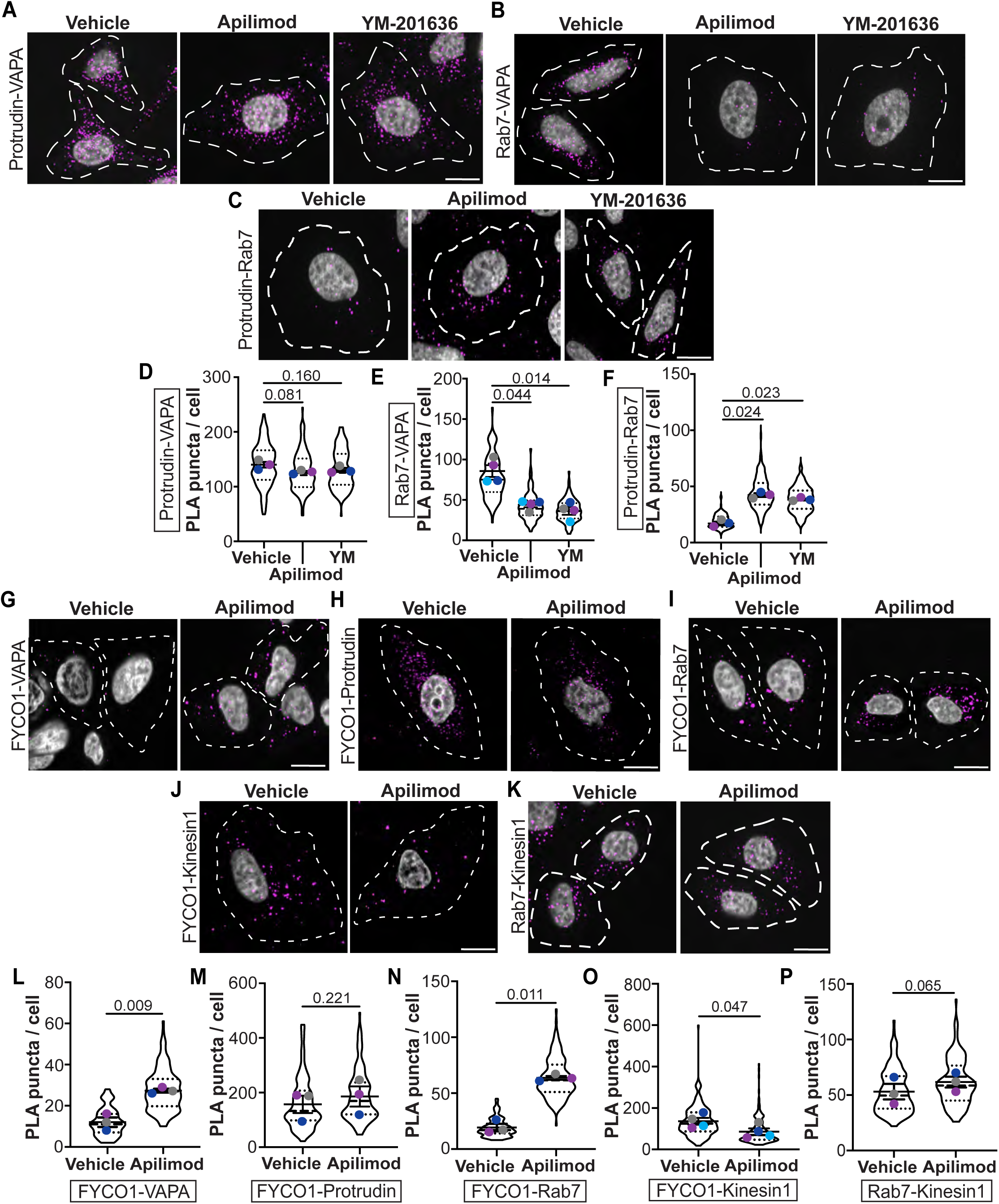
Loss of PIKfyve activity likely alters the molecular configuration of ER-lysosome contact sites. **(A-C)** Confocal images of HeLa cells treated with 0.02% DMSO (vehicle), 240 nM apilimod, or 1 µM YM-201636 for 2 h followed by Proximity Ligation Assay between protrudin and VapA (A), or Rab7 and VAPA (B), or Rab7 and protrudin (C). Scale bar = 15 µm. **(G-K)** Confocal images of HeLa cells treated with 0.02% DMSO (vehicle) or 240 nM apilimod for 2 h followed by Proximity Ligation Assay between FYCO1 and VAPA (G), or protrudin (H), or Rab7 (I), or kinesin-1 (J), and between Rab7 and kinesin-1 (K). Images were generated by maximum projection of z-stacks of the PLA signal (magenta); DAPI-stained nucleus is shown in gray. Negative controls where one of the primary antibodies was missing are shown in Sup. Fig.5A-H. Scale bar = 15 µm. **(D-F, L-P)** Quantification of the number of PLA dots per cell for each protein pair tested in A-C and G-K, as indicated. Experiments in D-F and K were repeated four independent times, while those in G-K were done three times. Data points were colour coded to match to respective experiment. Shown is the mean ± SEM. Data were analysed using an Ordinary one-way ANOVA and Dunnett’s multiple comparisons test (D-F) and paired Student’s t-test (L-P); p values are shown.

Protrudin-Rab7-VapA are proposed to load kinesin onto the Rab7-FYCO1 complex (Matsuzaki et al., 2011; Raiborg et al., 2015a; b; Pankiv et al., 2010). Moreover, PIKfyve and PtdIns(3,5)P_2_ may regulate kinesin-5 on precursor vesicles in neurons (Rizalar et al., 2023). Given our observations above and the collapse of lysosome motility and perinuclear accumulation, we next determined if PIKfyve inactivation altered the relative proximity of FYCO1 and kinesin-1 with these proteins. Unfortunately, we could not test the proximity between protrudin and kinesin-1 due to the lack of compatible antibodies. Collectively, there was an increase in proximity events between FYCO1 and Rab7 and VAPA, but no change in proximity events with protrudin (**Fig. 5G-I, 5L-N**). Importantly, there was a significant drop in FYCO1 and kinesin-1 proximity (**Fig. 5J, 5O**), which did not translate to altered Rab7 and kinesin-1 proximity (**FIG. 5K, 5P)**.

Overall, these data suggest that PIKfyve suppression reorganizes the conformation of the Rab7-VAPA-protrudin ER-lysosome contact site, whereby protrudin-Rab7 interaction is promoted at the expense of Rab7-VAPA proximity. Similarly, FYCO1 may be overly recruited to Rab7 and proximal to VAPA at the expense of kinesin-1 interaction, which may lead to reduced lysosome anterograde movement. Since PIKfyve inhibition concomitantly increases PtdIns(3)P while reducing PtdIns(3,5)P_2_ levels (Choy et al., 2018; Zolov et al., 2012) and protrudin and FYCO1 have a FYVE domain, we postulate that the excess PtdIns(3)P causes hyper-tethering onto lysosomal membranes. In fact, since protrudin is an ER transmembrane protein, we propose that hyper-stabilization of protrudin-Rab7 interaction via excess PtdIns(3)P alters the ER morphology.

### Protrudin and PtdIns(3)P are required for ER distortion upon PIKfyve inhibition

To test the hypothesis that protrudin drives ER distortion in the absence of PIKfyve activity, we quantified ER morphometrics in cells expressing ER-mScarletI, silenced for protrudin, and treated with apilimod. Surprisingly, protrudin silencing had no discernible effects on ER morphometrics in vehicle-treated cells (**Sup. Fig. S6A, B**; **Fig. 6A, 6C-D**), implying that protrudin does not have an essential role in defining basal ER morphology, or that redundant factors are at play. Similarly, protrudin silencing did not disturb lysosome volume or size (**Fig. 6B, 6E-F**). By contrast, protrudin silencing partially, but substantially offset the effects of apilimod on ER morphology (**Fig. 6A, 6C-D**), but not on lysosome metrics (**Fig. 6B, 6E-F**), indicating that protrudin is required for ER distortion in the absence of PIKfyve activity. We then treated ER-mNeonGreen-expressing cells with VPS34-In1, an inhibitor of VPS34, to block PtdIns(3)P and to determine if this lipid was also required for ER morphological changes during PIKfyve inactivation (**Fig. 6G**). Unlike apilimod alone, which changed ER morphometrics (**Fig. 6H, I**) and lysosome properties (**Fig. 6J, K**) relative to vehicle, VPS34-In1 alone and VPS34-In1 together with apilimod were not statistically different from vehicle for ER morphometrics (**Fig. 6H, I**) and lysosome properties (**Fig. 6J, K**; though lysosomes appeared somewhat enlarged). Together, these data intimate a role for protrudin and PtdIns(3)P in the reorganization of ER morphology triggered by PIKfyve inactivation.

**Figure 6.**
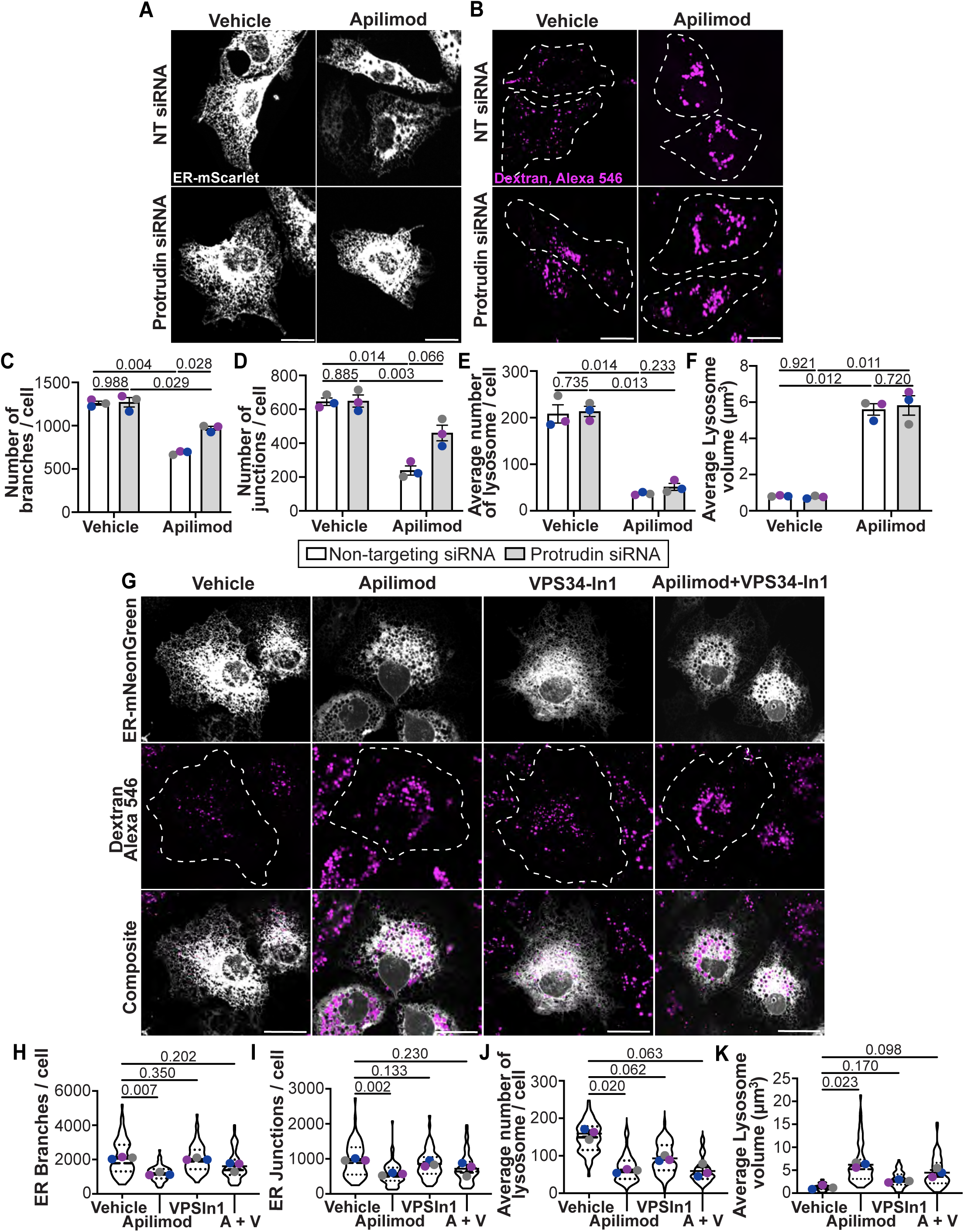
Protrudin and PtdIns(3)P are required for ER remodelling during PIKfyve inhibition. **(A)** Confocal images of U2OS cells co-transfected with non-targeting siRNA control or siRNA oligonucleotides against protrudin, and ER-mScarletI (gray), or **(B)** labeled with fluorescent dextran (magenta). **(C-F)** Quantitative analysis of ER and lysosomal morphology based on ER-mScarletI or dextran fluorescence in conditions shown (A): number of ER branches per cell (C), number of ER junctions per cell (D), average number of lysosome per cell (E), and average lysosome volume (F). **(G)** Confocal images of COS-7 cells transfected with ER-mNeonGreen (gray), labelled with fluorescent dextran (magenta), and then exposed to either 0.02% DMSO (vehicle), 240 nM apilimod, 1 µM VPS34-In1, or apilimod and VPS34-In1 for 2 h. Scale bar: full size = 20 µm **(H-K)** Quantification of ER morphology including number of ER branches per cell (H), number of ER junctions per cell (I), and quantification of average lysosome number per cell (J), and average lysosome volume (K). All experiments were repeated three independent times. Data points from matching independent experiments are colour coded. Shown are the mean ± SEM. Data were based on 20-25 transfected cells per condition per experiment. Data in (C-F) was analyzed using a two-way ANOVA and Tukey’s multiple comparisons test, and data in (H-K) was analyzed using a one-way ANOVA and Dunnett’s post-hoc test; p values are shown.

### Protrudin lipid tethering mechanisms are altered upon PIKfyve inhibition

We next expressed GFP-chimeras of wild-type (protrudin^wt^) and a FYVE4A-mutant protrudin (protrudin^FYVE4A^; (Raiborg et al., 2015a)) in cells expressing and/or silenced for endogenous protrudin (**Fig. 7A**). Protrudin^wt^-GFP and protrudin^FYVE4A^-GFP both exhibited ER-like distribution in the presence of (non-targeting) or the absence of endogenous protrudin (siRNA; **Fig. 7A**). Remarkably, cells expressing protrudin^wt^-GFP but inhibited for PIKfyve displayed intense protrudin^wt^-GFP ring structures that decorated both dextran-labelled and dextran-negative vacuoles (**Fig. 7A**). Since protrudin is an ER-transmembrane protein, this means that the ER membrane or regions of the ER membrane wrap around vacuoles induced in PIKfyve-disabled cells. This concept is supported by 3D reconstruction of protrudin^wt^-GFP, which extends and at least partially encloses dextran-loaded vacuoles (**Sup. Fig. S6C**). Importantly, the protrudin^FYVE4A^-GFP mutant failed to form these rings around dextran-loaded lysosomes in PIKfyve-inhibited cells (**Fig. 7A**), indicating that the protrudin-labelled ER membranes tightly encircle the enlarged lysosomes, likely in a PtdIns(3)P-dependent manner. Consistent with this, apilimod-treated cells displayed intense 2FYVE-RFP vacuoles that were reduced upon VPS34-In1 exposure, suggesting high PtdIns(3)P levels (**Fig. 7D**).

**Figure 7.**
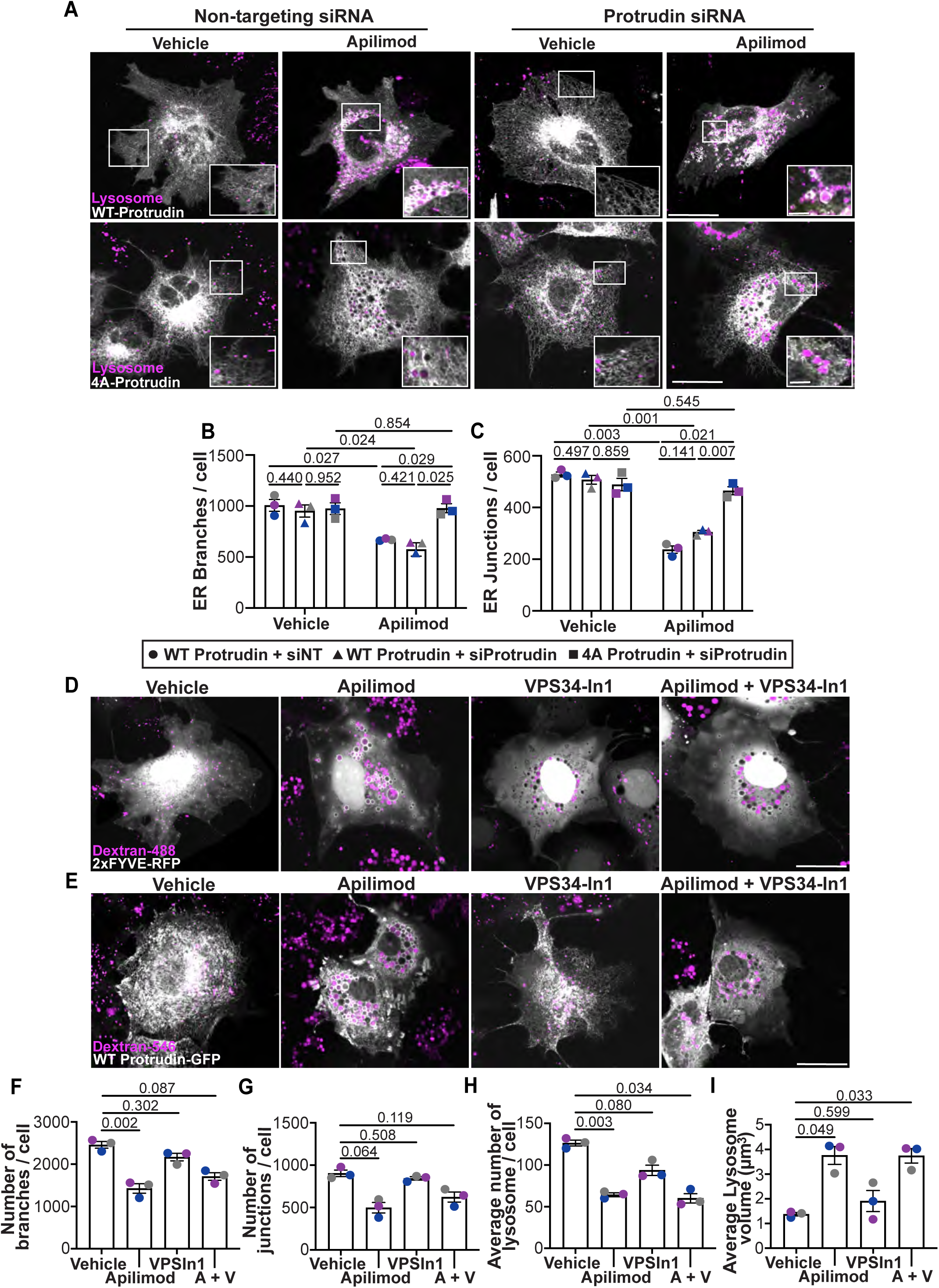
Protrudin hyper-tethers ER and lysosomes during PIKfyve inhibition to alter ER morphology. **(A)** Confocal images of HeLa cells co-transfected with non-targeting siRNA control or siRNA oligonucleotides against protrudin, and protrudin^wt^-GFP or protrudin^FYVE4A^-GFP (gray). Cells were labeled with fluorescent dextran (magenta) and then exposed to either 0.02% DMSO (vehicle) or 240 nM apilimod for 2 h. Scale bar: full size = 20 µm, zoom insert = 5 µm. **(B, C)** Number of protrudin branches per cell (B) and junctions per cell (C) based on protrudin^wt^-GFP or protrudin^FYVE4A^-GFP fluorescence. **(D,E)** Confocal images of COS7 cells transfected with 2FYVE-RFP (D; grayscale) or protrudin^wt^-GFP (E; grayscale), labelled with fluorescent dextran (magenta), and then exposed to either 0.02% DMSO (vehicle), 240 nM apilimod, 1 µM VPS34-In1, or apilimod and VPS34-In1 for 2 h. Scale bar: full size = 20 µm. **(F, G)** Quantification of ER morphology by number of ER branches per cell (F) and number of ER junctions per cell (G) in cells shown in E. **(H, I)** Quantification of lysosome number per cell (H) and lysosome volume (I) in cells shown in E. All experiments were repeated three independent times. Data points from matching independent experiments are colour coded. Shown are the mean ± SEM. Data are based on 20-25 transfected cells per condition per experiment. Data from (B-C) was analyzed using a three-way ANOVA and Tukey’s multiple comparisons test, and data from (F-I) was analysed using a one-way ANOVA and Dunnett’s post-hoc test; p values are shown.

We then determined if excess protrudin wrapping around vacuoles was at least partially responsible for the altered ER morphometrics in PIKfyve-arrested cells by skeletonizing the protrudin-GFP fluorescence. First, protrudin-GFP-based ER morphometrics was indistinguishable when cells were mock-silenced or silenced for endogenous protrudin (**Fig. 7B-C**, first two columns). Second, apilimod elicited dramatic changes to ER morphometrics in cells expressing wild-type protrudin-GFP, whether mock or silenced for endogenous protrudin (**Fig. 7B-C**, compare columns 1 and 3, and 2 and 4). We then compared the ER morphometrics by expressing protrudin^FYVE4A^-GFP in cells silenced for endogenous protrudin. In vehicle-treated cells, the ER morphology was like those expressing protrudin^wt^-GFP (**Fig. 7B-C**, compare columns 2 and 5). Yet, the ER changes driven by apilimod were annulled in cells expressing protrudin^FYVE4A^-GFP relative to protrudin^wt^-GFP (**Fig. 7B-C**, compare columns 4 and 6). Additionally, protrudin^wt^-GFP vacuoles were mostly eliminated when VPS34-In1 was co-incubated with apilimod (**Fig. 7E**), which partially recovered the ER morphology based on protrudin^wt^-GFP fluorescence (**Fig. 7F, G**), despite altered lysosome morphometrics (**Fig. 7H, I)**. Thus, a functional FYVE domain on protrudin and PtdIns(3)P are necessary for the apilimod-induced changes to ER morphometrics.

Since protrudin interacts with sculpting factors as well, we also assessed the distribution of Atlastin-1, CLIMP63, and Rnt4a. First, and as expected, apilimod changed the ER morphometrics based on overexpressed CLIMP63, Rtn4, and Atlastin (**Sup. Fig. S7A-F, H-I**). Secondly, apilimod-treated cells formed rings of Atlastin-1 (**Sup. Fig. S7G**) that were less prominent than protrudin-GFP (**Fig. 7**); CLIMP63 and Rtn4a rings were even less prominent, but visible (**Fig. 1A, B**). Together, it may be that the nanoscale organization of ER sculpting factors could further contribute to ER remodeling in PIKfyve-impaired cells.

### Excess protrudin lipid tethering during PIKfyve inhibition alters ER-lysosome contacts

We then explored if the absence of PtdIns(3)P or expression of protrudin^FYVE4A^-GFP rescued the relative proximity of protrudin-Rab7 and Rab7-VAPA in PIKfyve-defective cells. First, as a control, we note that protrudin silencing eliminated the number of proximity events between Rab7-protrudin PLA (**Fig. 8A, C)** and Rab7-VAPA PLA **(Fig. 8B, D),** the latter result suggesting protrudin cannot be removed without disturbing the contact sites containing Rab7 and VAPA. Third, we compared the proximity of these proteins in cells expressing protrudin^wt^-GFP or protrudin^FYVE4A^-GFP upon endogenous protrudin silencing **(Fig. 8E)**. As before, apilimod promoted Rab7-protrudin interactions in cells expressing endogenous (Fig. 8E, columns 1 and 4) and protrudin^wt^-GFP (**Fig. 8E**, columns 2 and 5). Importantly, there was no difference between vehicle and apilimod-treated cells expressing protrudin^FYVE4A^-GFP (**Fig. 8E**, columns 3 and 6) and a significant distinction between protrudin^wt^-GFP and protrudin^FYVE4A^-GFP with apilimod (**Fig. 8E**, columns 5 and 6).

**Figure 8.**
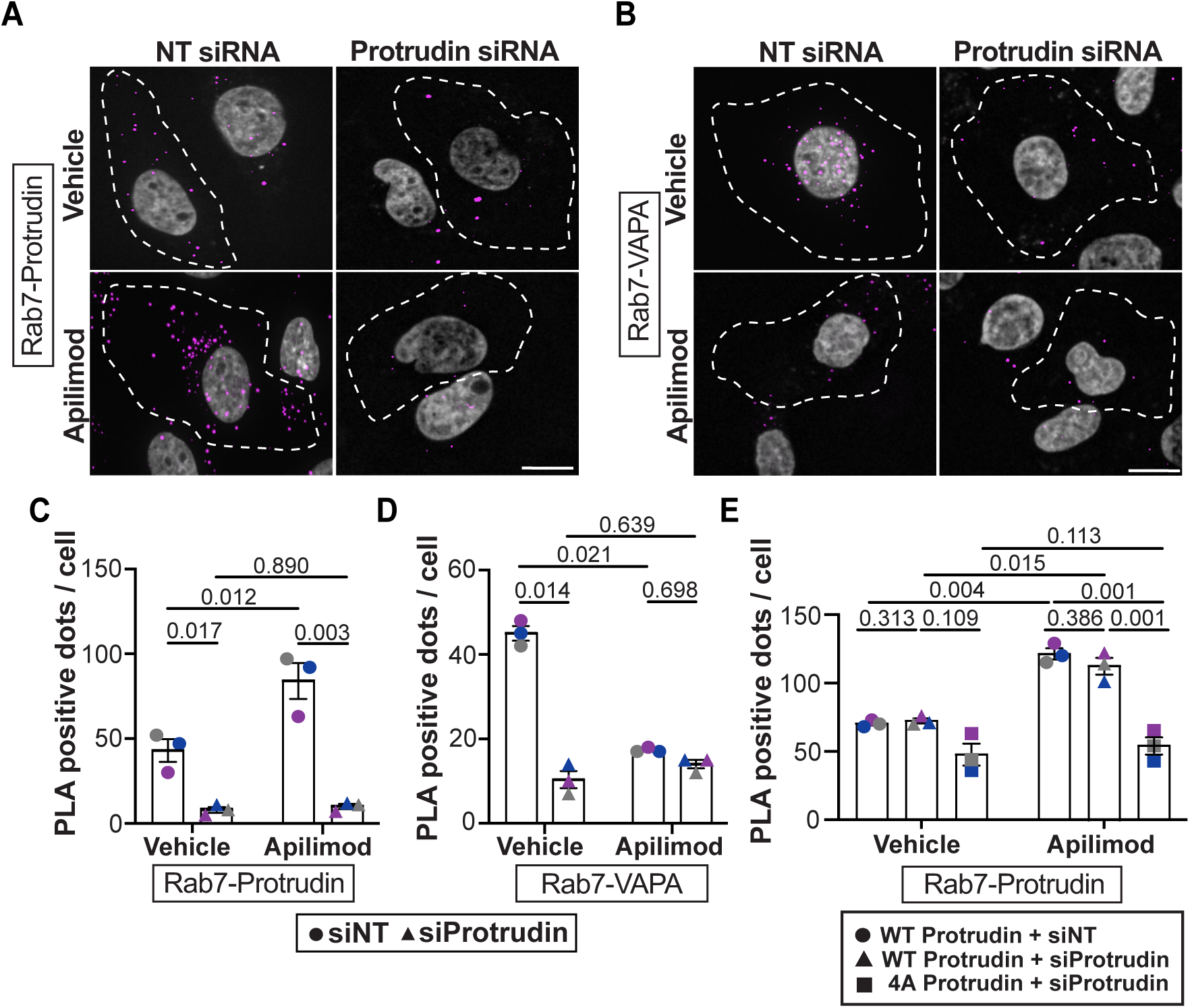
Protrudin and its FYVE domain are required for ER-lysosome changes induced by PIKfyve inactivation. **(A-B)** Confocal images of HeLa cells transfected with non-targeting siRNA control or siRNA oligonucleotides against protrudin and treated with 0.02% DMSO (vehicle) or 240 nM apilimod for 2 h followed by Proximity Ligation Assay between protrudin and Rab7 (A), or Rab7 and VAPA (B). Scale bar: full size = 20 µm. **(C-D)** Quantification of number of PLA dots between protrudin and Rab7 (C) and Rab7 and VAPA (D) in control silenced and protrudin-silenced cells with and without apilimod. **(E)** Quantification of PLA dots between Rab7 and protrudin in mock-silenced (siNT) or protrudin-silenced cells expressing protrudin^wt^-GFP or protrudin^FYVE4A^-GFP exposed to either vehicle or apilimod. All experiments were repeated three independent times and colour-code matched. Shown is the mean ± SEM. Data for (C-D) were analyzed using a two-way ANOVA and Tukey’s multiple comparison test, and data for (E) was analyzed using a three-way ANOVA and Tukey’s multiple comparisons test; p-values are shown.

Overall, we propose that PIKfyve inhibition leads to excess PtdIns(3)P levels, which inappropriately recruits and locks protrudin to lysosomes via its FYVE domain. This hyper-recruitment distorts the balance of ER-lysosome contact sites by shifting interactions toward Rab7-protrudin tethering and away from a conformation wherein Rab7-VapA are more proximal. The disproportionate lysosomal localization of protrudin also disrupts ER morphology.

### Enforced ER-lysosome contacts suffice to disrupt ER morphology

Our data suggest that PIKfyve inhibition accumulates PtdIns(3)P that hyper-tethers protrudin on lysosomes, disrupting the ER morphology. To test whether hyper-tethering of protrudin to lysosomes sufficed to disturb the ER architecture, we employed the FKBP-FRB inducible dimerization system by expressing iRFP-FRB-Rab7 and newly engineered FKBP-mCherry-protrudin^WT^ or FKBP-mCherry-protrudin^FYVE4A^ constructs. Without rapamycin, and based on their fluorescent signal, the ER morphometrics appeared normal for both WT and protruin^FYVE4A^ and no obvious wrapping of ER occurred around Rab7 puncta (**Fig. 9A).** In remarkable contrast to this, addition of rapamycin to dimerize the FKBP and FRB chimeric proteins, caused extreme ER wrapping around Rab7-positive lysosomes that strongly distorted ER morphometrics both for protrudin^WT^ and protrudin^FYVE4A^ (**Fig. 9A, 9B-D**, compare columns 1 and 2 with 3 and 4, respectively). Additionally, VPS34-In1, which could reverse changes to ER morphometrics by apilimod and GFP-protrudin (**Fig. 7E-G**), was impotent at rescuing the rapamycin-induced ER morphological changes and wrapping around lysosomes (**Fig. 9**). Overall, these data demonstrate that enforced ER-lysosome contact suffices to disturb ER morphology and bypass the need for the protrudin’s FYVE domain and PtdIns(3)P. In conclusion, this supports that the ER morphological changes we observed during PIKfyve inactivation are driven by excess PtdIns(3)P and hyper-tethering of protrudin to lysosomes.

**Figure 9.**
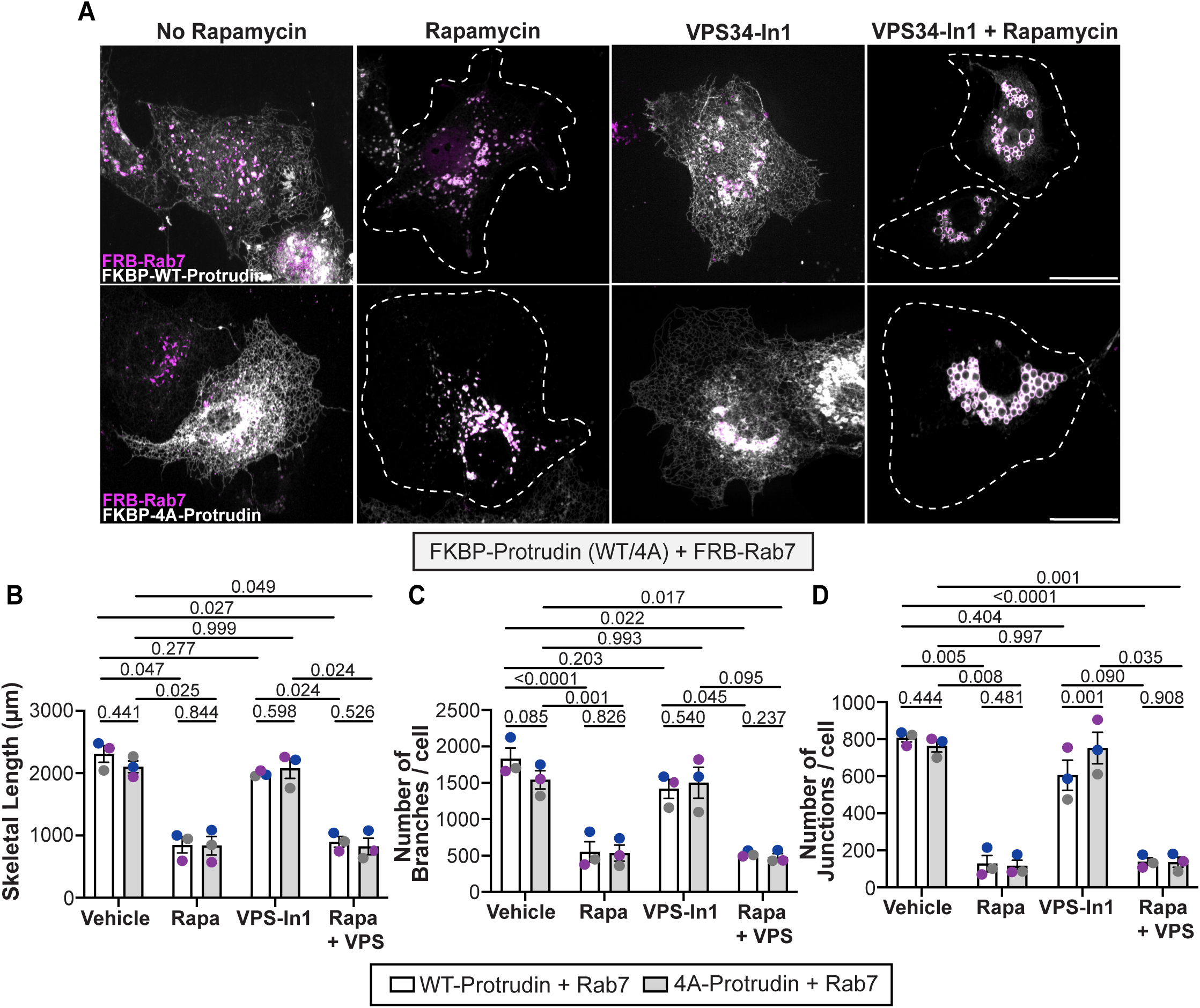
ER-lysosome hyper-tethering with inducible dimerization system suffices to distort the ER architecture. **(A)** Confocal images of COS-7 cells transiently transfected with iRFP-FRB-Rab7 (magenta) and FKBP-mCherry-protrudin^wt^ or FKBP-mCherry-protrudin^FYVE4A^ (gray). Cells were exposed to either no treatment, 100 nM rapamycin for 15 min, VPS34-In1 for 2 h, or VPS34-In1 and rapamycin combined. Scale bar: full size = 20 µm. **(B-D)** Quantification of protrudin-based ER morphology including skeletal length (B), number of branches per cells (C), and number of junctions per cell (D) in the indicated conditions. Experiments (B-D) were repeated three independent times and colour-code matched. Shown is the mean ± SEM. Data for (F-H) were analyzed using a two-way ANOVA and Tukey’s multiple comparisons test; p-values shown.

## Discussion

PIKfyve is a well-established regulator of the endolysosomal pathway (Choy et al., 2018; Bissig et al., 2017; Sbrissa et al., 2002; Ikonomov et al., 2001; Sharma et al., 2019). Here, we extend PIKfyve’s sphere of influence by revealing that PIKfyve activity affects ER morphology and dynamics – cells inhibited for PIKfyve activity present a less reticulate and dynamic ER, ostensibly driven in part by two mechanisms. First, the enlarged lysosomes become less motile, collapsing to the cell centre – as a corollary, we observed fewer ER hitchhiking events, which promotes ER remodelling and motility (Spits et al., 2021; Guimaraes et al., 2015; Langley et al., 2025; Lu et al., 2020). Second, the ER membrane appears “glued” to enlarged lysosomes by extensive protrudin-dependent binding, likely due to a surfeit of PtdIns(3)P (Choy et al., 2018; Zolov et al., 2012). Thus, we identified an important role for PIKfyve in modulating ER properties beyond those known for the endo-lysosomal system.

### PIKfyve modulation of lysosome and ER architecture

Inhibition of PIKfyve and PtdIns(3,5)P_2_ elicits lysosome coalescence (Choy et al., 2018; Bissig et al., 2017; Sharma et al., 2019). Currently, we do not know if this occurs due to dysregulation of fission factors in mammalian cells like the recently discovered HPO-27 (Li et al., 2024), and/or through regulation of ion signaling modulated by channels like TPCs, TRPLM1, and ClC7 (Leray et al., 2022; Dong et al., 2010; Wang et al., 2012), or through a different mechanism. Given that ER tubules contact and demarcate sites of mitochondrion and endosome fission (Rowland et al., 2014; Friedman et al., 2013, 2011), we sought to test if PIKfyve and ER-dependent fission were linked by overexpressing CLIMP63 and Rtn4a to respectively abate and promote ER tubule formation. We reasoned that promoting sheets, CLIMP63 overexpression would lead to enlarged lysosomes, mimicking PIKfyve arrest, while excess Rtn4a might rescue lysosome coalescence in PIKfyve-hindered cells by promoting ER tubules; however, we did not discern these predicted behaviours. While we observed effects on individual lysosome volume in CLIMP63/Rnt4a-modified cells, this was not accompanied by altered lysosome number, suggesting that the effects on lysosome size are not because of altered fusion-fission cycling, as is the case for PIKfyve (Choy et al., 2018; Sharma et al., 2019). Instead, we anticipate that processes such as modified biosynthesis and/or lipid transfer may drive these changes in lysosome size. It remains possible that CLIMP63 and Rtn4a overexpression are not ideal approaches to test the relationship between PIKfyve-driven processes and ER.

Conversely, disturbing PIKfyve greatly perturbed the ER architecture, which became less reticulated and more sheet-like, decorated with various spherical “voids” filled with enlarged lysosomes or endosomes. While some of these ER morphometric changes may reflect steric hindrance by the enlarged endosomes and lysosomes, our data point to a more elegant role for PIKfyve in modulating ER properties – by calibrating PtdIns(3)P levels to balance protrudin-dependent ER-lysosome contact sites. Supporting this model, ER morphometrics were partly rescued in PIKfyve-inhibited cells by *i)* VPS34-In1 co-administration with apilimod, *ii)* silencing of endogenous protrudin, and *iii)* re-expression of the FYVE4A-protrudin mutant, but not of the wild-type protrudin – all despite enlarged lysosomes. Convincingly, artificially clamping ER to lysosomes with rapamycin-inducible system sufficed to alter ER morphometrics even in the absence of PtdIns(3)P (**Fig. 9**). Thus, we propose that physical “glueing” of the ER membrane to lysosomes likely helps transform the ER architecture in PIKfyve-defective cells. However, physical obstruction by enlarged endosomes and lysosomes and nanoscale re-organization of ER morphogenic factors like CLIMP63, Atlastin-1, and reticulons may also contribute to ER remodeling in the absence of PIKfyve activity since these proteins interact with protrudin (Chang et al., 2013; Hashimoto et al., 2014).

### PIKfyve modulates lysosome and ER dynamics

A striking consequence of PIKfyve inhibition is the concurrent loss of lysosome and ER motility, which we propose are related processes. While microtubule morphometrics and dynamics appeared nearly normal in PIKfyve-inhibited cells, the enlarged lysosomes were perinuclear and less motile. While we do not know how PIKfyve modulates lysosome dynamics, an attractive mechanism is that PIKfyve tightly balances PtdIns(3)P and PtdIns(3,5)P_2_ levels in space and time to govern various PtdIns(3)P-interacting proteins such as FYCO1, RUFY3, RUFY4, protrudin, and SNX19, most of which modulate motor function (Pankiv et al., 2010; Saric et al., 2021; Raiborg et al., 2015a; Jongsma et al., 2024; Kumar et al., 2022; Keren-Kaplan et al., 2022; Jongsma et al., 2016, 2024). Remarkably, our observations on protrudin behaviour align well with previous work by Saric *et al. on* SNX19, another ER transmembrane protein that carries a PX domain that binds to PtdIns(3)P (Mas et al., 2014; Saric et al., 2021). These authors observed that SNX19 forms an ER-lysosome contact site that restricts endo-lysosome motility and that enlarged lysosomes in PIKfyve inhibited cells were extensively decorated with SNX19 and were less motile (Saric et al., 2021). Collectively, it is intriguing to consider that PIKfyve is not just important to synthesize PtdIns(3,5)P_2_, but crucial to tightly modulate the levels and location of PtdIns(3)P to coordinate these proteins. Temptingly, this may explain the existence of the conserved PIKfyve complex, which holds together the PIKfyve and the antagonistic Fig4 lipid phosphatase (Botelho et al., 2008; Jin et al., 2008; Sbrissa et al., 2007; Botelho, 2009) – to exquisitely switch between PtdIns(3)P and PtdIns(3,5)P_2_ at spatially-restricted sites like membrane contacts. This phosphoinositide-based switch may interface with the previously observed role of repeated ER-endosome contacts and microtubule-associated proteins in swapping motor activity on endosomes (Raiborg et al., 2015a; Jongsma et al., 2024), and regulation of ions like Ca^2+^(Li et al., 2016)a.

As for loss of ER dynamism, we propose that this is partly due to declined lysosome motility, and consequently, reduced ER hitchhiking, which is a major driver of ER remodeling (Spits et al., 2021; Lu et al., 2020). However, ER hitchhiking also depends on the formation of lysosome-ER contact sites. Based on our observations, ER-lysosome contact sites not only form, but may in fact undergo hyper-tethering during PIKfyve arrest as visualized for protrudin by us and for SNX19 by *Saric et al*, both of which indicate that ER membrane subdomains envelope enlarged lysosomes and related organelles. This implies that bidirectional transport of lysosomes and ER may depend on regulated conformational or compositional changes to these contacts, which consequently alter kinesin and/or dynein-mediated activity, discussed next.

### PIKfyve and regulation of ER-lysosome contact sites

Protrudin is thought to load kinesin-1 onto FYCO1 on lysosomes (Raiborg et al., 2015a). To do this, protrudin binds to PtdIns(3)P and Rab7 on lysosomes and VAPA on the ER membrane (Raiborg et al., 2015a). Nonetheless, there is little known about the structural organization of these proteins at these contact sites – for example, we do not know if all interactions are concurrent, sequential, or if subject to conformational changes. Since PIKfyve inhibition did not lead to a wholesale loss or gain in proximity events between protrudin, VAPA, Rab7, FYCO1, and kinesin-1, this suggests that PIKfyve activity aids in a conformational or selective compositional change at these sites. For example, protrudin-Rab7 and Rab7-FYCO1 proximity increased, while VAPA-Rab7 and FYCO1-kinesin1 decreased in PIKfyve-inactive cells. Importantly, apilimod-enhanced Rab7-protrudin proximity events depended on a functioning FYVE domain of protrudin. Collectively, we propose a model whereby PIKfyve converts PtdIns(3)P to PtdIns(3,5)P_2_ to transiently promote protrudin dissociation from lysosomal membranes and elicit a conformational change that helps modulate kinesin-1 loading and/or activity. Of course, PtdIns(3,5)P_2_ itself may have a regulatory role in this process, but we do not know the targets of this lipid in this context, if any.

## Conclusion

Collectively, our results suggest that PIKfyve supports ER network integrity by governing lysosome dynamics, distribution, and ER-lysosome contact sites. In the absence of PIKfyve, swollen and immobile lysosomes gain excess PtdIns(3)P that hyper-tether ER-lysosomes via protrudin, trapping these in a non-productive conformation that alters ER architecture and hitchhiking. These results illuminate a previously underappreciated aspect of PIKfyve function and provide mechanistic insight into how lysosomal dysfunction can drive architectural and dynamic changes of neighbouring organelles. Moreover, the PIKfyve pathway is linked to neurological diseases and is a potential therapeutic target against certain cancers and viral infections (Chow et al., 2007; Cai et al., 2013; Babu et al., 2024; Cheng et al., 2025; Bao et al., 2023; Kang et al., 2020). The physiological link between the PIKfyve pathway and these diseases is generally thought to occur through changes to the endo-lysosomal pathway. However, given our findings, mutations in the PIKfyve pathway that lead to disease may reflect altered ER function as well, particularly in highly secretory or polarized cells where proper ER architecture and dynamics are critical.

## Methods and Materials

### Cell culture

COS-7 and HeLa cell strains were cultured in Dulbecco’s Modified Eagle Medium (DMEM; Wisent, St. Bruno, QC) and U2OS-wt cells were cultured in 50% Dulbecco’s Modified Eagle Medium / 50% F-12 with L-glutamine and HEPES buffer (F-12/DMEM; Gibco, New York, USA) supplemented with 10% heat-inactivated fetal bovine serum (FBS; Wisent) and 1% penicillin-streptomycin (10,000 U/mL) (P/S; Gibco, Burlington, ON). All cells were authenticated, checked for contamination and cultured at 37°C with 5% CO_2_.

### Plasmids, siRNA gene silencing, and transfection

Plasmids obtained from Addgene were as follows: ER-mNeonGreen (#137804) and ER-mScarletI (#137805) were described in (Chertkova et al., 2020); pHAGE2 GFP-Rtn4a (#86684) and pHAGE2 GFP-Atlastin1 (#86678) were described in (Wang et al., 2016); mCherry-CLIMP63 (#136293, (Shibata et al., 2008)); eGFP-KIF5B (#172203, (Guardia et al., 2016)); pmRFP-C3::Rab5 (#14437, (Vonderheit and Helenius, 2005)); iRFP-FRB-Rab7 (#51613; (Hammond et al., 2014)); pcDNA3.1-mCherry (#128744), pcDNA3-eGFP (#13031), and pEGFP-N1::human EB1 (#39299). The plasmid encoding 2xFYVE-RFP was a gift from Dr. Amra Saric at the Hospital for Sick Children. We note that ER-mNeonGreen and ER-mScarletI are fusions of the mNeonGreen and mScarletI fluorescent proteins to the KDEL ER-retention signal, respectively (Chertkova et al., 2020). Plasmids encoding protrudin^WT^-GFP, protrudin^FYVE4A^-GFP were a kind gift from Dr. Camilla Raiborg (Oslo University Hospital) and was previously described (Raiborg et al., 2015a). Plasmids encoding protrudin^WT^ or protrudin^FYVE4A^ mutant fused to FKBP-mCherry were generated by GenScript (Piscataway, NJ, USA). Protrudin sequences were cloned at the C-terminus of the FKBP-mCherry construct. The FKBP-mCherry backbone (Addgene plasmid #108122) was originally generated by Gerry Hammond. Protrudin DNA sequences were kindly provided by Eva Maria Wenzel (Institute for Cancer Research, Oslo, Norway). Transient transfections were performed with FuGENE HD (Promega, Madison, WI) following the manufacturer’s instructions, using a 3:1 DNA to transfection reagent ratio. The transfection mixture was replaced with fresh medium after 4-6 h and cells were used 24 h post-transfection.

Silencing RNA oligonucleotides were transfected into HeLa and U2OS cells with Lipofectamine RNAiMAX Transfection Reagent (Invitrogen, Carlsbad, CA) following the manufacturer’s instructions. For protrudin silencing, one round of knockdown was performed using 25 nM ON-TARGETplus ZFYVE27 (the standard gene name for protrudin) individual siRNA (Horizon Discovery, Dharmacon; J-016349-12-0010) or ON-TARGETplus non-targeting control (Horizon Discovery, Dharmacon) over 24 h. For Vac14 silencing, one round of knockdown was performed using 25 nM ON-TARGETplus Vac14 siRNA individual siRNA 1 (Horizon Discovery, Dharmacon; Oligo 8, J-015729-08-005) and siRNA 2 (Horizon Discovery, Dharmacon; Oligo 5, J-015729-05-005) or ON-TARGETplus non-targeting control (Horizon Discovery, Dharmacon) over 48 h.

### Organelle labelling

Lysosomes in COS-7 were labelled with 320 µg/mL Alexa-Fluor 488-, 546- or 647-conjugated dextran (ThermoFisher), while lysosomes in HeLa and U2OS-wt cells were labelled with 150 µg/mL of fluorescent dextran by incubating cells in cell-specific complete medium for 1 h at 37°C with 5% CO_2_, followed by washing in phosphate-buffered saline (PBS) and replenishing with fresh complete medium for 1 h before live-cell imaging (Choy et al., 2018; Ohkuma and Poole, 1978; Ferris et al., 1987). Since this method likely labels a spectrum of late endosomes, lysosomes and endolysosomes, the term *lysosomes* reflects this range of organelles in this work. mCherry-Rab5 labels early endosomes, specifically (Vonderheit and Helenius, 2005; Bucci et al., 1992; Chavrier et al., 1990). ER-mNeonGreen and ER-ScarletI were used to label bulk ER (Chertkova et al., 2020).

### Pharmacological treatments

PIKfyve inhibition was accomplished by treating cells with 80-240 nM apilimod (Selleck Chemicals LLC, Houston, TX) or 1 µM YM-201636 (Cayman Chemical Company, Ann Arbor, MI) for 1-2 h – specific concentrations and times are indicated in each figure legend. Vps34 Class III PI 3-kinase was arrested by treating cells with 1 µM VPS34-In1 (Chemical Company, Ann Arbor, MI) for 2 h, and when used in combination with apilimod, VPS34-In1 was co-incubated for 2 h.

### Immunofluorescence

For immunofluorescence, following treatments, cells were fixed with 4% paraformaldehyde (PFA; v/v) for 15 min. After washing with PBS, residual PFA was quenched with 100 mM glycine for 10 min, then cells were blocked and permeabilized with 0.1% (v/v) Triton X-100 for 5 min. Cells were then subjected to mouse monoclonal (DM1A) antibody against ⍺-tubulin (1:100; #T9026; Sigma-Aldrich), rabbit monoclonal (D9F52) antibody against Rab7 (1:100; #9367), mouse monoclonal (E9O7E) antibody against Rab7 (1:100; #95746), mouse monoclonal (D4O1S) antibody against LAMP1 (1:100; #15665) and rabbit monoclonal (D2D11) antibody against LAMP1 (1:200; #9091; all from Cell Signaling Technology). Dylight-conjugated secondary donkey antibodies against rabbit and mouse (1:1000; Novus Biologicals Canada) were used.

### Proximity ligation assays

PLA was performed with Duolink In Situ Red Starter Kit Mouse/Rabbit (Sigma Aldrich, DUO92101) according to the manufacturer’s instructions. Briefly, HeLa cells were seeded, fixed with 4% PFA for 15 min, quenched with 100 mM glycine for 10 min and permeabilized with 0.1% Triton X-100 for 5 min – all at RT respectively. Cells were co-incubated with one mouse and one rabbit antibody from the following; rabbit monoclonal (D9F52) antibody against Rab7 (1:100; #9367; Cell Signaling Technology), mouse monoclonal (E9O7E) antibody against Rab7 (1:100; #95746; Cell Signaling Technology), mouse monoclonal (4C12) antibody against VAPA (1:100; #293278; Santa Cruz Biotechnology), rabbit polyclonal antibody against ZFYVE27 (1:100; 12680-1-AP; Proteintech), rabbit monoclonal (E3H4J) antibody against KIF5B (1:250; #47610; Cell Signaling Technology), rabbit polyclonal antibody against FYCO1 (1:250; 25730-1-AP; Proteintech), rabbit and mouse monoclonal (5G4) antibody against GFP (1:500; #55494; Cell Signaling Technology), followed by incubation with PLA probes. After probe incubation, cells were washed and incubated with ligation mixtures to promote DNA ligation. Next, amplification mixtures were added to cells for Rolling Circle Amplification (RCA) and hybridization of fluorescently labelled oligonucleotides to occur. A PLA mounting media containing DAPI was added to washed cells when mounted. PLA signals from cells were detected using confocal microscopy.

### Fluorescence microscopy

Confocal images were obtained using a Quorum Diskovery spinning disc confocal system (Quorum Technologies, Inc., Guelph, ON), a Quorum ReSCAN confocal system (Quorum Technologies, In., Guelph, ON) and a Olympus CSU spinning disc confocal system (Evident Scientific, Tokyo, Japan). The Diskovery microscope consisted of an inverted fluorescence microscope (DMi8; Leica) equipped with an Andor Zyla 4.2 Megapixel sCMOS camera (Oxford Instruments, Belfast, UK), a 63x oil immersion objective (1.4 NA), and standard excitation and emission filter sets and lasers were used for all fluorophores. The microscope system was controlled by the MetaMorph acquisition software (Molecular Devices, LLC, San Jose, CA, USA). The ReSCAN microscope consisted of an inverted fluorescence microscope (DMi8 Automated; Leica) equipped with a ORCA-Flash V3 digital sCMOS camera (Hamamatsu, Shizuoka Prefecture, Japan) , a 63x oil immersion objective (0.6 to 1.4 NA) , and standard excitation and emission filter sets and lasers were used for all fluorophores. The microscope system was controlled by the Volocity acquisition software (Quorum Technologies, In., Guelph, ON). The Olympus spinning disc microscope consisted of an inverted fluorescence microscope (IX83, Olympus) equipped with a ORCA-Flash4.0 V3 sCMOS camera (Hamamatsu, Shizuoka Prefecture, Japan), a 60x oil immersion objective (UPLAPO OHR; NA 1.49; refractive index 1.518; working distance 100 µm), and standard excitation and emission filter sets and lasers were used for all fluorophores. The microscope system was controlled by the CellSens acquisition software (Evident Scientific, Tokyo, Japan). For all live-cell imaging, cells were maintained in DMEM free of phenol red in a Chamlide microscope-mounted chamber at 37°C and 5% CO_2._ For z-stack imaging, 20-50 images were acquired with an interplanar distance of 0.2 to 0.6 µm, where origin was set to the centre of the cell. For timelapse obtained for ER and lysosome dynamics and EB1-GFP dynamics, frames were acquired every 0.25 s for 1 min. For timelapse obtained for ER-hitchhiking, frames were acquired every 0.3 s for 1 min.

### ER and lysosome morphology analysis

Image analysis was conducted using Volocity and Fiji software. ER morphology analysis was conducted on Volocity using an ER protocol designed to provide skeletal length measurements through the thinning (or skeletonizing) of the ER until only a single-voxel-wide-line remains (Arganda-Carreras et al., 2010; Schindelin et al., 2012). The protocol then counts the number of voxels along the path and provides information such as skeletal length, curvature, branching and fragmentation. Further ER morphology and fluorescent ER-protein distribution, analyses were completed using a semi-automated junction analysis plug-in for Fiji provided by Spits *et al (2021)* from the Leiden University Medical Centre in the Netherlands. This plug-in was based on the analyze skeleton feature on Fiji where an ER mask is initially created and the skeleton representation is analyzed for the number of junctions and branches (Spits et al., 2021). Lysosome morphology analysis was done on Volocity using a “Measure Objects” protocol designed to provide voxel/pixel count, volume and area, and total number measurements for the entire cell (Hipolito et al., 2019; Saffi et al., 2021).

### ER and lysosome dynamics analysis

ER dynamics were analyzed using a membrane displacement analysis macro for Fiji also provided by *Spits et al (2021)* that analyzed mobile ER fractions based on #pixels/frame ranging from 0-9. This method is largely based on optical flow analysis which compares corresponding areas in two frames and generates a map that depicts the frame-to-frame movement of specific image areas. The extent of movement is reflected by the change in pixel color intensity between each frame (Spits et al., 2021). Seven separate 60-pixel x 60-pixel regions of interest were randomly selected in a clockwise manner targeting both peripheral and perinuclear ER for one single cell per field of view for seven cells and the mobile ER fractions were determined. We then binned movement into three categories of change (0-3 #pixels/frame), (4-7 #pixels/frame) and (8-9 #pixels/frame) to define low-, intermediate-, and rapid-movement.

Lysosome dynamics were analyzed using Single Particle Tracking analysis for Fiji, which analyzed lysosome dynamics upon PIKfyve inhibition using time-lapse images acquired every 0.25 s for 1 min. Default settings were kept with the LoG detector (Laplacian of Gaussian). In vehicle-treated cells, the diameter of the lysosome was set to 1.5 µm with a quality threshold of 500. A uniform color was selected for color spots. The simple LAP Tracker was selected with the linking max distance set to 7 µm, the gap-closing distance set to 5 µm, and the gap closing max frame set to 2 µm. A track index was generated, and the following parameters were extracted: total distance travelled, mean speed, and overall track displacement. In PIKfyve inhibited cells, the same process was followed with the exception that the diameter of the lysosome was set to 2 µm with a quality threshold of 250; and both the linking max distance and gap-closing max distance were set to 5 µm. Lysosome motility was also shown using a “Reslice” protocol in Fiji to create Kymographs between conditions, where a segmented line was drawn along the x-axis of the cell randomly, with a line width of 2 pixels; generating a space-time projection displaying spatial position (x-axis) versus time (y-axis).

The frequency of ER-hitchhiking events upon PIKfyve inhibition was analyzed using Time-lapse analysis acquired every 0.25 s for 1 min. Cells post-treatment were analyzed to determine the number of ER-hitchhiking events for multiple, randomized (100 µm^2^) regions of interest (ROI). To quantify the number of events, Fiji software was utilized to select ROIs based on available ER periphery with co-migrating lysosomes. A positive hitchhiking event was identified if the lysosome localized with the tip of a growing ER tubule, initiated movement, co-travelled with the ER tubule, and “dropped it off” at another independent ER tubule. Hitchhiking events were tracked and identified qualitatively by eye.

### Lysosome distribution analysis

To determine the change in lysosome distribution, Shell analysis on Fiji software was utilized. Using the Freehand Selection Tool on Fiji, the cell was outlined to create a “total cell” ROI. This was followed by using the Enlarge Tool on Fiji to create a shell by shrinking the “total cell” ROI by 30-50 pixels to create a second layer of shell. This process was repeated to create a third and fourth layer of shell. The raw integrated density was calculated for each shell independently to determine the sum of the values of pixel intensities in the respective shell. The raw integrated density of the innermost shell as a fraction of the “total cell” ROI was calculated and compared to the outermost shell as a fraction of the “total cell” ROI.

### 3D Image Reconstruction

Three-dimensional (3D) image reconstruction was performed using Imaris Software (V9.5, Bitplane, Oxford Instruments). Raw z-stack images (plane thickness: 0.2 µm) were acquired using a ReSCAN Confocal Microscope (previously explained) and imported into Imaris for analysis. Prior to reconstruction, image stacks were subjected to background subtraction preprocessing, with a rolling ball background subtraction of 50 pixels applied across all images and conditions using Fiji software. Within Imaris, 3D volumes were generated using the Surpass Mode, and surfaces were rendered using the Surface or Volume modules, with thresholds set manually based on signal intensity. Reconstruction parameters, including smoothing and thresholding method were consistently applied across all samples to ensure comparability. For visualization, coloring was applied to distinguished channels representing lysosomes (magenta) and protrudin (gray).

### Recombinant protein expression and purification

GST-fused RILP-C33 (truncated mutant) and GST-fused empty vectors (ampicillin resistant) were expressed in BL21 competent cells, grown at 37 °C and shaking at 200 rpm over-night . Cells were cultured until the OD_600_ reached 0.6. After treatment with 1 µM isopropyl-ß-D-thiogalactoside (IPTG) for 4 h at 30 °C and shaking at 200 rpm, the induced recombinant GST-RILP-C33 and GST proteins were centrifuged at 4 °C for 30 minutes at 5000 xg, with the supernatant discarded and pellet resuspended in 1X bacterial lysis buffer (50 mM Tris, 150 mM NaCl, 10 mM β-mercaptoethanol, pH 7.5 with protease inhibitor cocktail (Sigma-Aldrich). The bacterial lysates were then homogenized using a EmulsiFlex-C3 pressure homogenizer (AVESTIN, Canada). Once homogenized, the recombinant GST-RILP-C33 and GST- lysates were centrifuged at 4 °C for 30 minutes at 5000 xg, pelleted, and the supernatant was incubated with 50% glutathione-Sepharose 4B bead resuspension made following manufacturer’s instructions (Millipore Sigma, G317-0756-01) at a dilution of 1:50 (beads to bacterial lysate supernatant, respectively) for 2 h at 4 °C with gentle shaking. The samples were then centrifuged at 4 °C for 5 minutes at 5000 xg, pelleted, supernatant discarded, and pellet resuspended in 1X bacterial lysis buffer. The samples were analyzed by western blotting to confirm protein expression and purification.

### Affinity precipitation of GTP-bound Rab7

HeLa cells were pre-treated with either vehicle control (DMSO), apilimod, or YM-201636 and lysed in 700 µL modified- 1X RIPA Buffer (25 mM Tris HCl, 150 mM NaCl, 1% TritonX-100, 1% Na deoxychlorate, pH 7.4 with protease inhibitor cocktail (Sigma-Aldrich)). Cells were then incubated on ice for 30 minutes, collected into clean microcentrifuge tubes and centrifuged at 4°C for 5 minutes at 3,000 xg. Prior to centrifugation, 10% of each lysate sample was collected reserved as an input control. These input samples were also centrifuged at 4°C for 5 minutes at 3,000 xg and subsequently mixed with a 1:1 ratio of 2X Laemmli Sample Buffer (0.5 M Tris, pH 6.8, glycerol and 10% sodium dodecyl sulfate (SDS), 1:10 β-mercaptoethanol). Recombinant RILP-C33 and GST bound to glutathione-Sepharose 4B beads previously explained were mixed with HeLa cell lysates at a dilution of 1:14 (beads to cell lysate, respectively). These mixtures were incubated for 2 h at 4 °C with gentle shaking. The samples were then centrifuged at 4°C for 5 minutes at 500 xg, with remaining supernatant discarded. The sample pellets were eluted using modified-1X RIPA Buffer (1:2 ratio of sample to buffer, respectively). The supernatant from each wash was moved to a clean microcentrifuge tube and a 1:1 ratio of 2X Laemmli Sample was added to the samples, boiled at 96°C for 10 minutes and resolved via SDS-PAGE.

### SDS-PAGE and Western blotting

Following treatment, whole cell lysates were prepared in 200 µL 2x Laemmli Sample Buffer (0.5 M Tris, pH 6.8, glycerol and 10% SDS) supplemented with a protease and phosphatase inhibitor cocktail (Complete 1x protease inhibitor (Sigma-Aldrich), Halt phosphatase inhibitor cocktail (ThermoFisher Scientific)). Lysates were heated at 95°C for 10 min, and 10% b-mercaptoethanol and 5% bromophenol blue were added to the cell lysates. Proteins were resolved by Tris-Glycine SDS-PAGE and transferred on to a polyvinylidene difluoride (PVDF) membrane. The PVDF membrane was blocked for 1 h at RT in blocking buffer composed of 5% BSA in 1x final Tris-buffered saline-Tween (TBS-T; 20 nM Tris, 150 mM NaCl, and 0.1% Tween-20) After blocking, membranes were washed and incubated with primary antibody (at designated dilution) overnight at 4°C. Primary antibodies used for western blotting include: rabbit polyclonal antibody against Vac14 (1:1000, SAB4200074, Sigma Aldrich), rabbit polyclonal antibody against protrudin (1:1000, 12680-1-AP, Proteintech), rabbit monoclonal (14C10) antibody against GDPH (1:5000, #2118S, Cell Signaling Technology), rabbit monoclonal (DE3C6) antibody against clathrin (1:5000, #4796s, Cell Signaling Technology), rabbit monoclonal (D95F2) antibody against Rab7 (1:1000, #9367S, Cell Signaling Technology) and rabbit monoclonal antibody against beta-actin (1:5000, #4967, Cell Signaling Technology). After primary incubation, membranes were washed and then subjected to secondary antibody (at designated dilution) incubation for 1 h at RT. After incubation with secondary antibody, membranes were washed and imaged using the Chemi-Doc Imaging System (Bio-Rad). Western blot signals were analyzed and quantified using ImageLab 6.1 (Bio-Rad). Band intensity was quantified by signal integration in an area corresponding to the appropriate band.

### FKBP-FRB Inducible Dimerization

For inducible FKBP-FRB dimerization experiments, COS-7 cells were seeded at a density of 2.0 x 10^4^ cells/mL and cultured overnight. For baseline ER morphology analysis (in the absence of FKBP-FRB heterodimerization), cells were transfected with GFP-protrudin^WT.^ Following transfection, cells were washed with PBS and replenished with DMEM. Lysosomes were labelled using Alexa Fluor 546-Dextran, followed by treatment with either 0.02% DMSO (vehicle) or 240 nM apilimod treatment for 2 h. For inducible FKBP-FRB dimerization experiments, COS-7 cells were co-transfected with either protrudin^WT^ or protrudin^FYVE4A^ FKBP-mCherry constructs and iRFP-FRB-Rab7. Following transfection, cells were washed with PBS and replenished with DMEM. Cells were then treated with 1 µM VPS34-In1 for 2 h, or with 100 nM Rapamycin for 15 min, or rapamycin and VPS34-IN1 prior to imaging. Live-cell imaging was performed with z-stacks acquired (plane thickness = 0.2 µm).

### Statistical Analysis

All experiments were performed at least three independent times with the specific total number indicated in the respective figure legends along with sample size. The mean and measure of variation such as standard deviation (STD) or standard error of the mean (SEM) are indicated. When shown, violin plots indicate range, followed dotted lines to display 25-75% percentile, the dashed line indicates median of the sampled populations. Sample means of independent experiments are shown and colour coded to match the independent experiment across conditions. All experimental conditions were statistically analyzed with the actual tests disclosed in the respective figure legend. Unless data was normalized, sampling was assumed to be normally distributed. As such, data was typically assessed with paired Student’s t-test when comparing two populations, with a repeated-measures one-way ANOVA for three or more samples with one parameter, repeated-measures two-way ANOVA when two parameters were considered, or repeated-measures three-way ANOVA when three parameters were considered. The Geisser-Greenhouse correction was applied. Post-hoc tests were employed based on assumptions and comparisons as recommended by GraphPad Prism and disclosed in the respective figure legends when comparing multiple conditions in non-normalized controls. ANOVA tests were coupled to Dunnett’s Multiple Comparison Test for one-way analysis and Tukey’s multiple comparison test for two-way analysis, to analyze pairwise conditions. Here, we disclose all p values for greater transparency, while we employ the standard p<0.05 to support a statistical difference between populations, we view p<0.1 as indicative of a difference and p<0.01 as very high confidence of a difference.

## Data availability statement

All data and original tools built in this study will be available upon request.

## Acknowledgements

We would like to thank Dr. Camilla Raiborg (Oslo University Hospital) for kindly providing the plasmids encoding the GFP fusion of the wild-type and of the FYVE4A mutant of protrudin. We also thank Kimberly Lau at the Sick Kids Imaging Facility for advice on Imaris and 3D reconstruction.

## Abbreviations

Arl8b: ADP-ribosylation factor like 8b
Atg18: autophagy-related protein 18
ClC7: chlorine exchange transporter 7
CLIMP63: 63 kDa cytoskeleton linking protein
EB1: end-binding protein 1
ER: endoplasmic reticulum
FFAT: two phenylalanine’s (FF) in an acidic tract
Fig4: factor-Induced gene 4
FKBP: FK506-binding protein
FRB: FKBP rapamycin binding
FYCO1: FYVE and coiled-coil domain autophagy adaptor 1
FYVE: fab1, YOTB, Vac1 and EEA1 binding domain
FYVE4A: fab1, YOTB, Vac1 and EEA1 binding domain mutant
GFP: green fluorescent protein
GST: glutathione s-transferase
GTPase: guanine triphosphatase
KIF5B: kinesin-1 heavy chain isoform 5B
LAMP1: lysosomal associate membrane protein 1
LCR: low complexing region
mCherry/mCh: monomeric cherry, red fluorescent protein
MCS: membrane contact site
mScarletI: basic red fluorescent protein
PI(3,5)P_2_: phosphatidylinositol 3,5-bisphosphate
PI(3)P: phosphatidylinositol 3-phosphate
PIKfyve: phosphoinositide FYVE-type zinc finger containing
PIP: phosphoinositide
PtdIns: phosphatidylinositol
PLA: proximity ligation assay
PX Domain: phox homology domain
Rab5: ras-related protein-5
Rab7: ras-related protein-7
RFP: red fluorescent protein
RILP: rab interacting lysosomal protein
RILPC33: rab interacting lysosomal protein, c terminal 33aa fragment
Rtn4a: reticulon 4A
RUFY3: RUN and FYVE domain containing 3
RUFY4: RUN and FYVE domain containing 4
siRNA: small interfering RNA
SnxA: sorting nexin A
Snx19: sorting nexin 19
TAC: tip attachment complex
TPC: two pore channel
TRPML1: transient receptor potential channel for mucolipin-1
VAPA: vesicle associated membrane protein-associated protein A
VAPB: vesicle associated membrane protein-associated protein B
VPS34: vacuolar sorting protein 34
VPS34 In1: vacuolar sorting protein 34 inhibitor-1
2XFYVE: tandem FYVE domain.

## Supplementary Figure Legend

**Supplementary Figure S1. ER morphology and dynamics are altered upon PIKfyve inhibition using YM-201636. (A)** Confocal images of COS-7 cells transiently transfected with ER-mNeonGreen (gray) and labelled with Dextran, Alexa-647 (magenta). Cells were exposed to either 0.02% DMSO (vehicle) or 1 µM YM-201636 for 2 h. **(B-E)** Quantification of ER morphology including ER skeletal length (B), number of ER junctions per cell (C), number of branches per cell (D), and average branch length (E). **(F)** Representative time-lapse confocal images of COS-7 cells transiently transfected with ER-mNeonGreen (gray). Skeletonize function can be applied to trace ER morphology. Zoom inserts show ER redistribution over 40 s; white arrowheads exemplify morphological changes. Scale bar: full size = 20 µm, zoom insert = 5 µm. **(G-I)** Quantification of membrane displacement analysis categorized by low movement (G), intermediate movement (H), and rapid movement (I). Experiments (B-E) were repeated four independent times; Experiments (G-I) were repeated six independent times. Data points from matching independent experiments are colour coded. Shown in the mean ± SEM. Data from B through E are based on 15-20 transfected cells per condition per experiment. Data from G through I are based on 7 randomized ROIs for 7-10 transfected cells, per condition per experiment. Data were analyzed using a two-tailed Student’s t-test and p values are shown.

**Supplementary Figure S2. Voids in ER architecture are filled with vacuolated endosomes and lysosomes upon PIKfyve inhibition. (A-B)** Confocal images of COS-7 cells labeled with Dextran, Alexa Fluor 647 (magenta) and transiently expressing ER-mNeonGreen (gray) or mCherry (cyan) (A) or mRFP-Rab5 (cyan) (B). Cells were exposed to either 0.01% DMSO or 80 nM apilimod for 2 h. Scale bar: full size = 20 µm, zoom insert = 5 µm. **(C-D)** Quantification of Mander’s Analysis done on the colocalization between Dextra, Alexa 647 and mRFP-Rab5. All experiments were repeated three independent times. Data points from matching independent experiments are colour coded and shown as mean ± SEM from 30-50 transfected cells per condition per experiment. Data were analyzed using a two-tailed Students t-test and p values are shown.

**Supplementary Figure S3. Vac14 silencing disturbs ER morphology. (A)** Representative Western blot of Vac14 from U2OS cells transfected with non-targeting siRNA control (siNT) or one of two siRNA oligonucleotides against Vac14. GAPDH was used as a loading control. **(B)** Quantitative analysis displaying Vac14 protein levels normalized to GADPH after Vac14 silencing. **(C)** Confocal images of U2OS cells co-transfected with either non-targeting siRNA or siRNA oligonucleotides against Vac14 and ER-mScarletI (gray) followed by an immunofluorescence staining to LAMP1 (magenta). Scale bar: full size = 20 µm, zoom insert = 5 µm. **(D-G)** Quantitative analysis of ER morphology in control or Vac14-silenced cells using skeletal length (D), number of branches per cell (E), number of junctions per cell (F), and average branch length (G). All experiments were repeated three independent times. Data points from matching independent experiments are colour coded and shown as mean ± STD (B) and mean ± SEM (D-G). Data from (D-G) were obtained from 20-25 transfected cells per condition per experiment. Data were analyzed using a one-way ANOVA and Dunnett’s multiple comparisons test.

**Supplemental Figure S4. PIKfyve inhibition does not significantly affect microtubule architecture and dynamics. (A)** Confocal images of HeLa cells immunostained for ⍺-tubulin (gray) and LAMP1 (magenta) treated with 0.02% DMSO (vehicle), 240 nM apilimod or 1 µM YM-201636 for 2 h. Scale bar = 20 µm **(B-D)** Quantification of microtubule morphology including number of branches per cell (B), number of junctions per cell (C), and average branch length (D). For B through D, all experiments were repeated four independent times. **(E)** Confocal images of COS-7 cells transiently transfected with EB1-GFP displayed as Rainbow RBG LUT and treated as in A. Scale bar: full size = 20 µm, zoom insert = 5 µm. **(F-H)** Semi-quantification of microtubule dynamics including percentage of cells with >10 EB1-puncta (F), percentage of cells with mobile EB1-puncta (G), and percentage of cells with EB1-puncta moving away from the nucleus (H). For F-H, all experiments were repeated three independent times. For B-D and F-H, data was analyzed using an Ordinary one-way ANOVA test and Dunnett’s multiple comparisons test.

**Supplementary Figure 5. Rab7 GTPase status appears unaltered in PIKfyve-inhibited cells. (A-H)** Control confocal images of HeLa cells treated with one individual primary antibody only for each PLA test for protrudin and VAPA (A), Rab7 and VAPA (B), Rab7 and protrudin (C), FYCO1 and VAPA (D), FYCO1 and protrudin (E), FYCO1 and Rab7 (F), Rab7 and kinesin-1 (G) and FYCO1 and kinesin-1 (H). Scale bar= 15 µm. **(I)** HeLa cells immuno-stained for LAMP1 (magenta) and Rab7 (gray) following treatment with 0.02% DMSO (vehicle) and 240 nM apilimod for 2 h. Scale bar= 20 µm. **(J-K)** Quantitative analysis displaying the average Mander’s Coefficient for the fraction of Rab7 colocalized with LAMP1 (J), and fraction of LAMP1 colocalized with Rab7 (K). **(L)** Representative Western blots of affinity purification of Rab7-GTP using GST-RILPC33, displaying total Rab7 levels from input samples and GTP-Rab7 levels recovered with the GST negative control and GST-RILPC33 from cells exposed to 0.02% vehicle (DMSO), 240 nM apilimod or 1 µM YM-201636 for 2 h. **(M)** Quantitative analysis of affinity precipitated GTP-Rab7 from GST negative control and GST-RILPC33. GST-control and GST-RILPC33 were normalized first to input levels of β-actin and then Rab7. All experiments were repeated three independent times. Data points from matching independent experiments are colour coded. Data shown in J through K are shown as the mean ± SEM and analyzed using a two-tailed Student’s t-test. Data from M is shown as the mean ± STD and was analyzed using a two-way ANOVA and Tukey’s multiple comparisons test; p values are shown.

**Supplementary Figure 6. Three-dimensional reconstruction of protrudin-positive ER. (A)** Representative Western blots of non-targeting siRNA control or siRNA oligonucleotides against protrudin displaying protrudin and clathrin levels. **(B)** Quantitative analysis displaying protrudin protein levels upon transfection with siNT or siProtrudin after normalized to loading control, clathrin. **(C)** Three-dimensional reconstruction of HeLa cells co-transfected with non-targeting siRNA control or siRNA oligonucleotides against protrudin, and expressing protrudin^wt^-GFP or protrudin^FYVE4A^-GFP (gray). Cells were labeled with Dextran, Alexa 546 (magenta) and then exposed to either 0.02% DMSO (vehicle) or 240 nM apilimod for 2 h. Scale bar: full frame = 20 µm, zoom insert = 0.7 µm. All experiments were repeated three independent times. Data points from matching independent experiments are colour coded. Shown in the mean ± STD. Data was analyzed using a two-tailed Student’s t-test and p values are shown.

**Supplementary Figure 7. PIKfyve inhibition leads to global ER changes across different resident proteins. (A-F)** Quantitative analysis of COS7 cells transfected with mCherry-CLIMP63 (A-C, as shown in Fig. 1A) or GFP-Rtn4a (D-F, as in Fig. 1B) upon treatment with either 0.02% DMSO (vehicle) or 80 nM apilimod for 2 h. ER morphometrics were analyzed including skeletal length (A, D), number of branches per cell (B, E) and number of junctions per cell (C, F). **(G)** Confocal images of COS7 cells co-transfected with GFP-Atlastin1 and GFP-protrudin^wt^, followed by treatment with either 0.02% DMSO (vehicle) or 80 nM apilimod for 2 h. Scale bar = 20 µm. **(H-I)** Quantitative analysis of ER morphometrics using Atlastin-1 fluorescence using number of branches per cell (H) and number of junctions per cell (I). Experiments from (A-F) were repeated four independent times; experiments from (H-I) were repeated three independent times. Data points from matching independent experiments are colour coded. Shown is the mean ± SEM. Data was analyzed using a two-tailed Student’s t-test and p values are shown.

## Supplementary Video Legend

**Supplemental Video 1. ER motility is retained upon treatment with vehicle control (DMSO).** COS-7 cells were transiently transfected with ER-mNeonGreen (gray) and treated with 0.02% vehicle (DMSO) for 2 h. The ER network undergoes continuous remodeling through tubule extension, retraction, and branching, reflecting its highly dynamic and interconnected nature. Time-lapse images were acquired at a rate of 4 frame/s for 1 minute. Still frames from Video 1 are displayed in Figure 3A. Scale bar: 20 µm.

**Supplemental Video 2. ER motility is reduced upon acute treatment with PIKfyve inhibitor Apilimod.** COS-7 cells were transiently transfected with ER-mNeonGreen (gray) and treated with 240 nM apilimod for 2 h. Time-lapse images were acquired at a rate of 4 frame/s for 1 minute. Still frames from Video 2 are displayed in Figure 3A. Scale bar: 20 µm.

**Supplemental Video 3. ER motility is reduced upon acute treatment with PIKfyve inhibitor YM-201636.** COS-7 cells were transiently transfected with ER-mNeonGreen (gray) and treated with 1 µM YM-201636 for 2 h. Time-lapse images were acquired at a rate of 4 frame/s for 1 minute. Still frames from Video 3 are displayed in Supplemental Figure 2A. Scale bar: 20 µm.

**Supplemental Video 4. EB1-GFP dynamics under vehicle condition conditions.** COS-7 cells were transiently transfected with EB1-GFP displayed as a rainbow LUT scale, where red is the highest intensity level and blue is the lowest intensity level. Cells were treated with 0.02% vehicle (DMSO) for 2 h. EB1-GFP comets mark the plus-ends of growing microtubules, enabling visualization of dynamic microtubules. Time-lapse images were acquired at a rate of 4 frame/s for 1 minute. Still frames from Video 4 are displayed in Figure 4E. Scale bar: 20 µm.

**Supplemental Video 5. EB1-GFP dynamics appear unaltered upon PIKfyve inhibition using apilimod.** COS-7 cells were transiently transfected with EB1-GFP displayed as a rainbow LUT scale, where red is the highest intensity level and blue is the lowest intensity level. Time-lapse images were acquired at a rate of 4 frame/s for 1 minute. Still frames from Video 5 are displayed in Figure 4E. Scale bar: 20 µm.

**Supplemental Video 6. Microtubule EB1 dynamics remain unaffected by YM-201636 treatment.** COS-7 cells were transiently transfected with EB1-GFP displayed as a rainbow LUT scale, where red is the highest intensity level and blue is the lowest intensity level. Time-lapse images were acquired at a rate of 4 frame/s for 1 minute. Still frames from Video 6 are displayed in Figure 4E. Scale bar: 20 µm.

**Supplemental Video 7. Normal ER-movement via organelle hitchhiking in vehicle-treated cells.** COS-7 cells were transiently transfected with ER-mNeonGreen (gray) and labeled with Dextran, Alexa 647 (magenta) followed by treatment with 0.02% vehicle (DMSO) for 2 h. Positive hitchhiking events were scored based on ER movement driven by physical association with a motile dextran labelled lysosome. Timelapse images were acquired at a rate of 3.3 frames/s for 1 minute. Still frames from Video 7 are displayed in Figure 4J. Scale bar: 20 µm.

**Supplemental Video 8. ER-hitchhiking frequency drops upon PIKfyve inhibition.** COS-7 cells were transiently transfected with ER-mNeonGreen (gray) and labeled with Dextran, Alexa 647 (magenta) followed by treatment with 240 nM apilimod for 2 h. Positive hitchhiking events were scored based on ER movement driven by physical association with a motile dextran labelled lysosome. Timelapse images were acquired at a rate of 3.3 frames/s for 1 minute. Still frames from Video 7 are displayed in Figure 4J. Scale bar: 20 µm.

