## Supplementary figures and images for "PIKfyve influences inter-organelle contacts with lysosomes to modulate the endoplasmic reticulum"

### Supplemental Figure S1

SUPPLEMENTAL FIGURE S1.

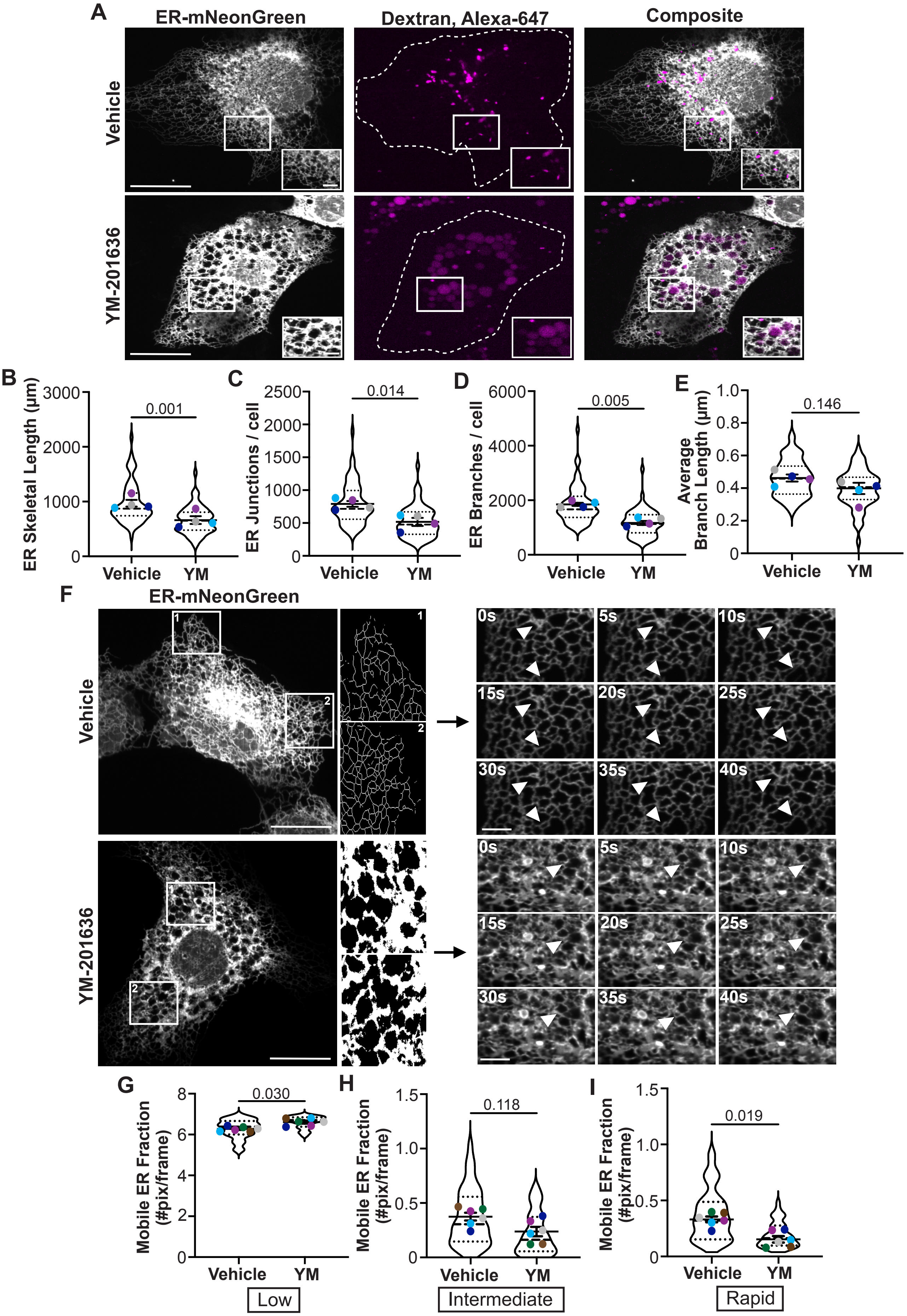

### Supplemental Figure S2

SUPPLEMENTAL FIGURE S2.

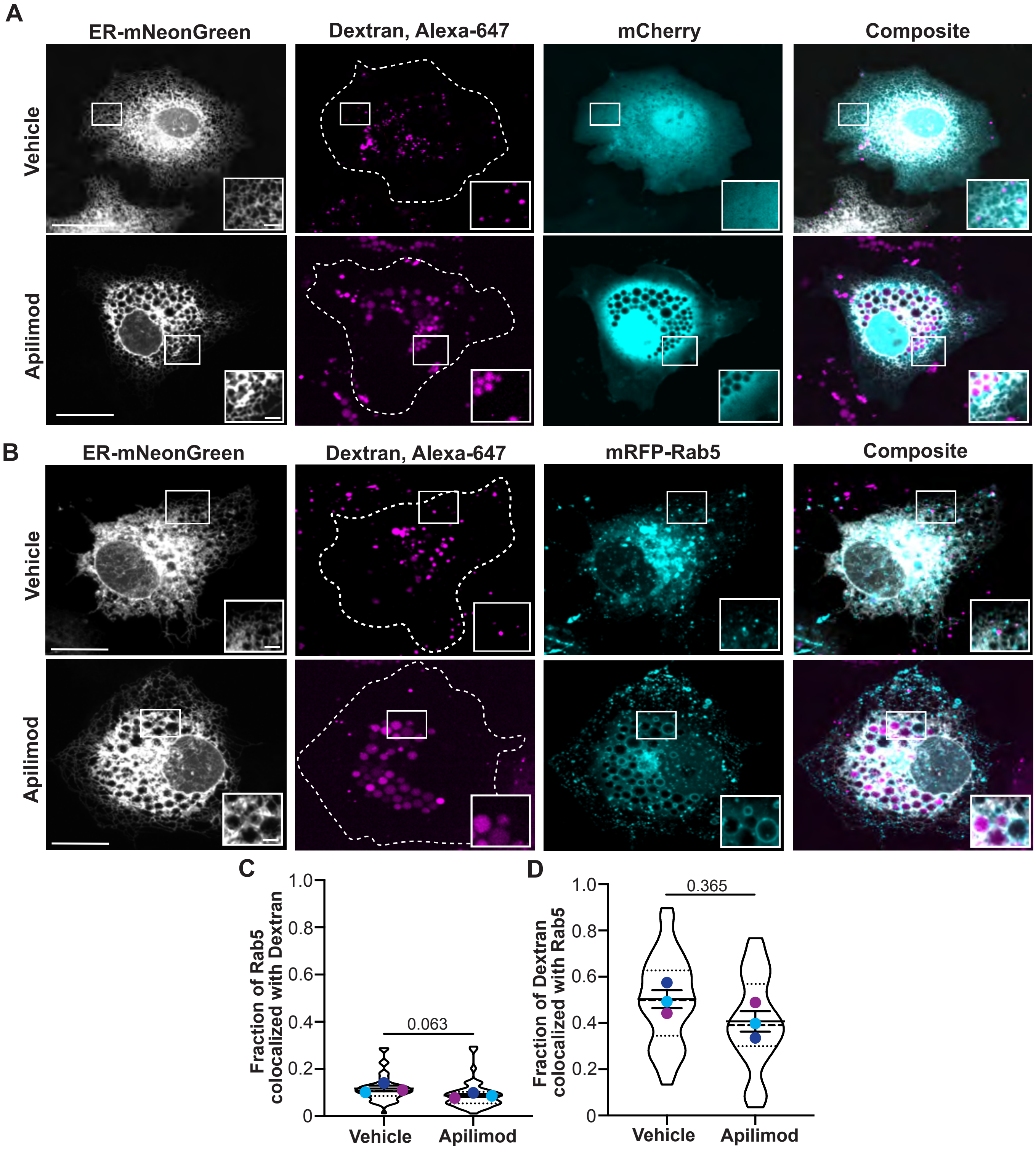

### Supplemental Figure S3

SUPPLEMENTAL FIGURE S3.

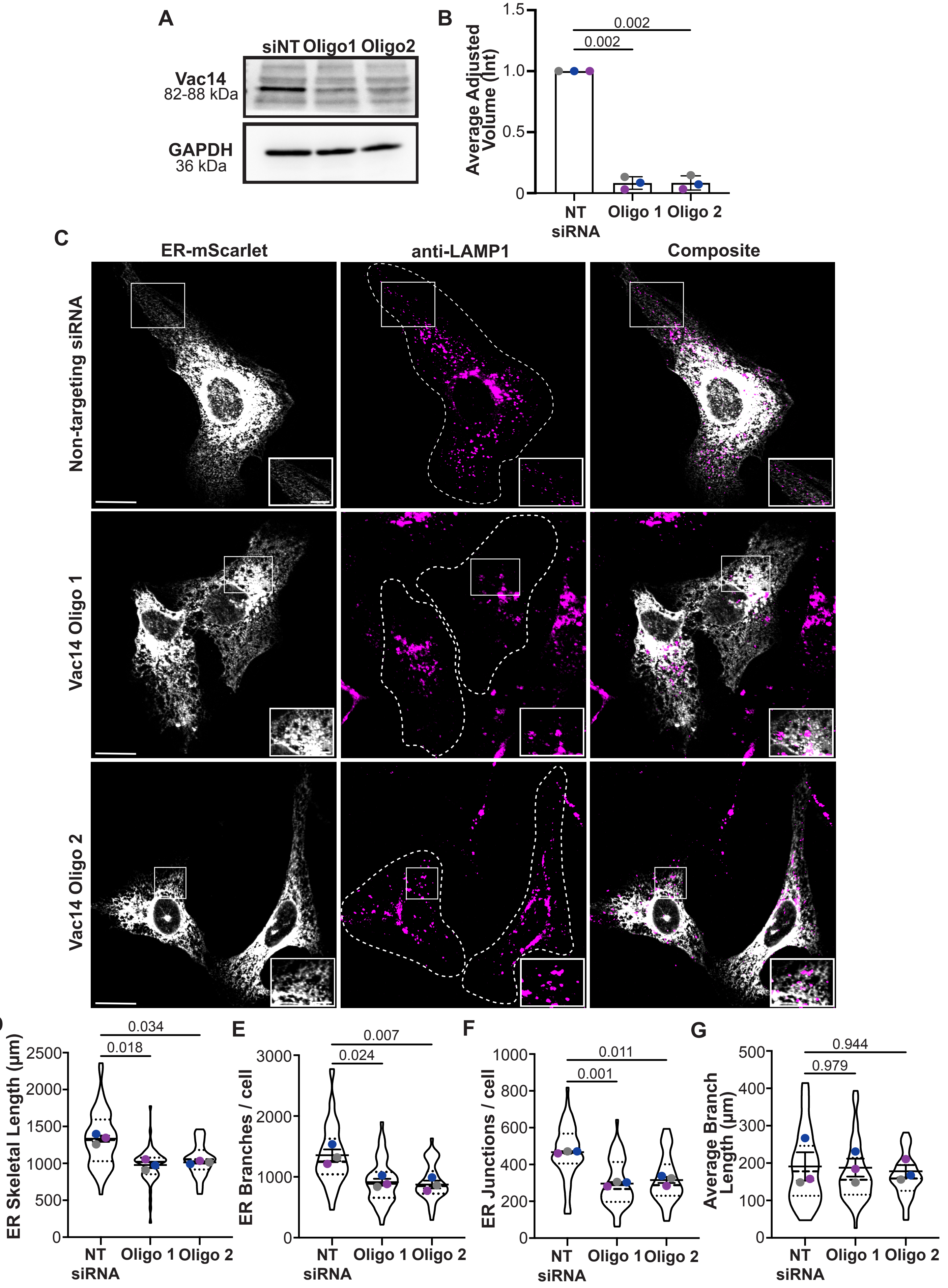

### Supplemental Figure S4

SUPPLEMENTAL FIGURE S4.

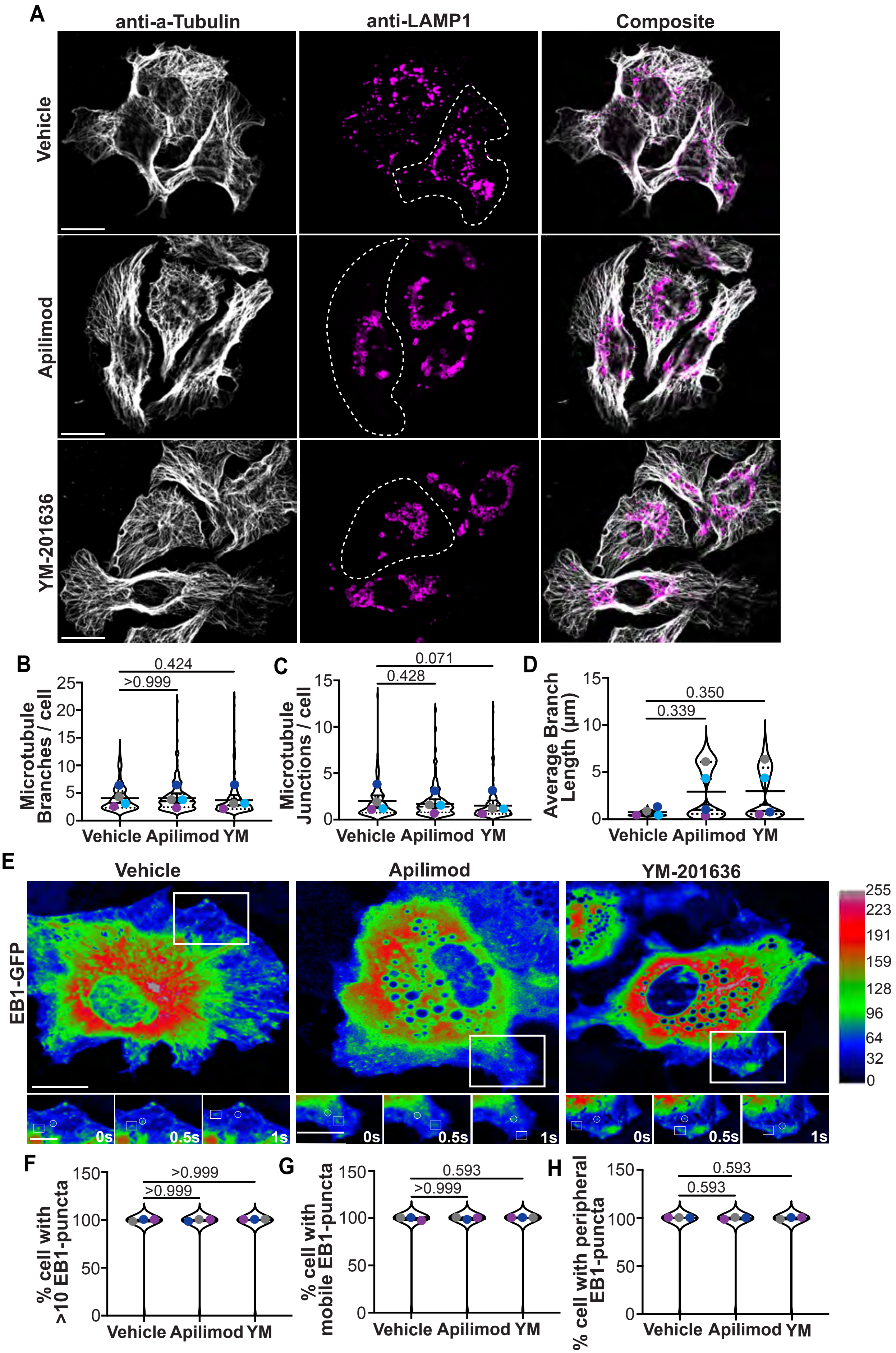

### Supplemental Figure S5

SUPPLEMENTAL FIGURE 5.

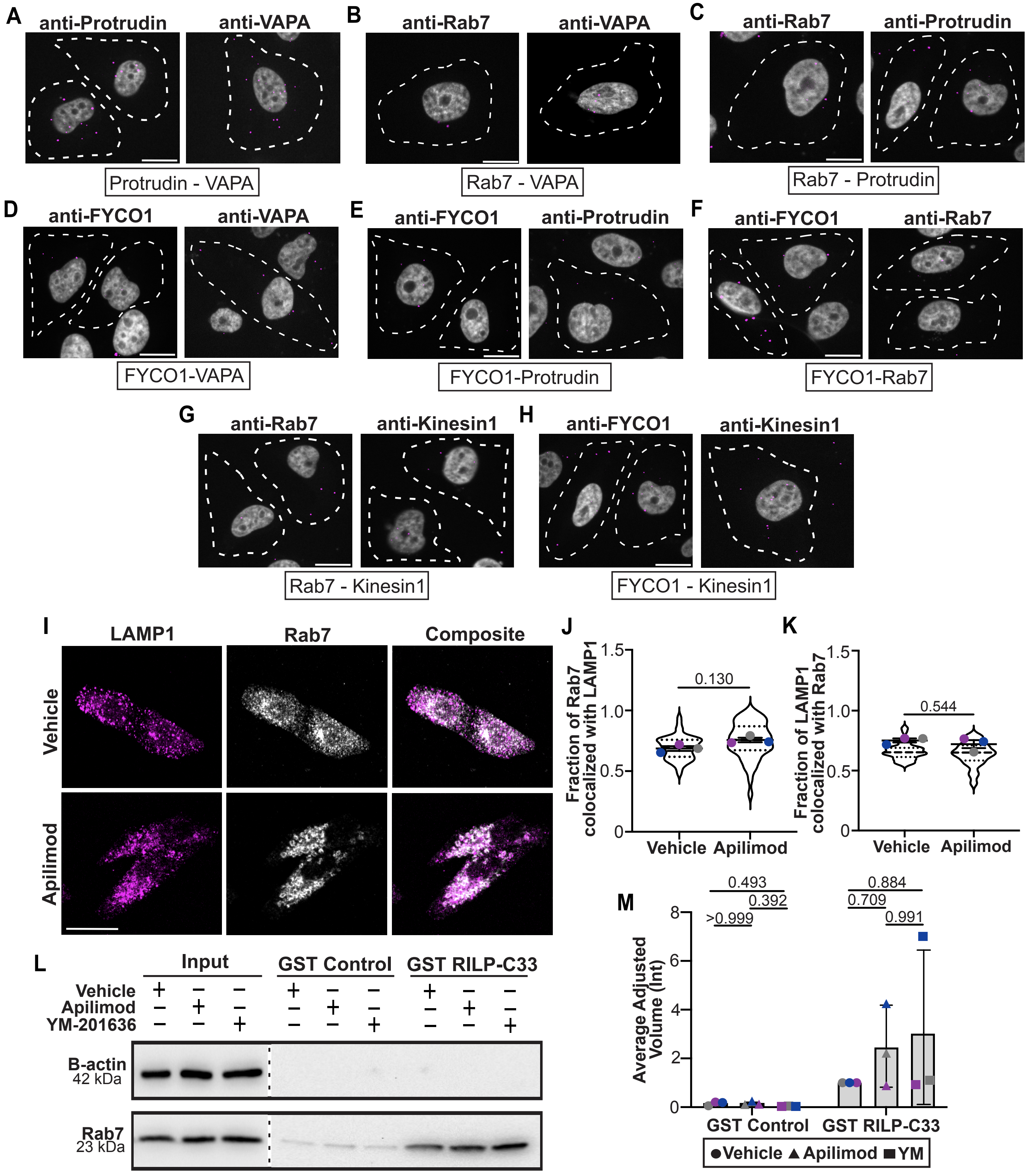

### Supplemental Figure S6

SUPPLEMENTAL FIGURE S6.

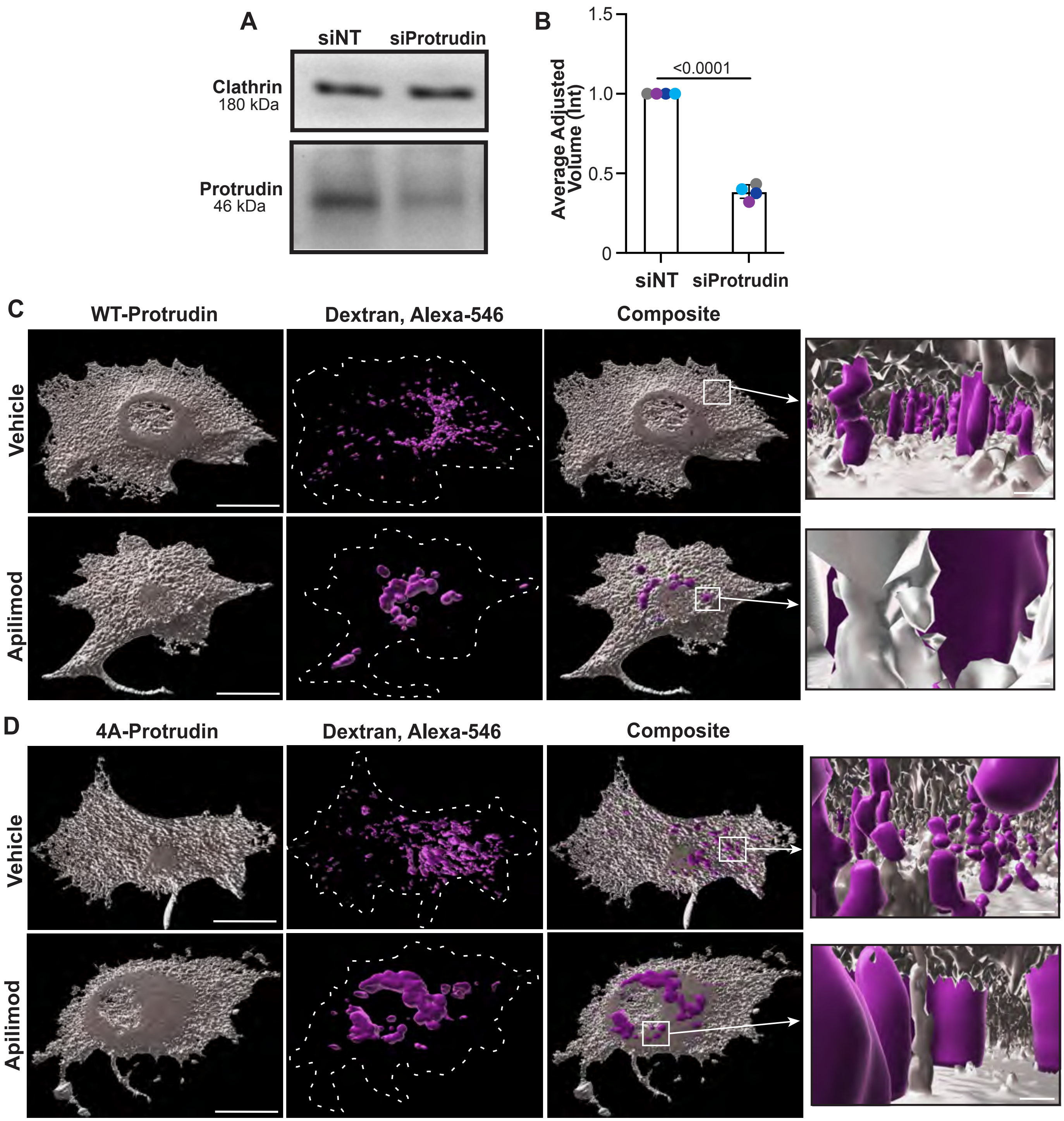

### Supplemental Figure S7

SUPPLEMENTAL FIGURE S7.

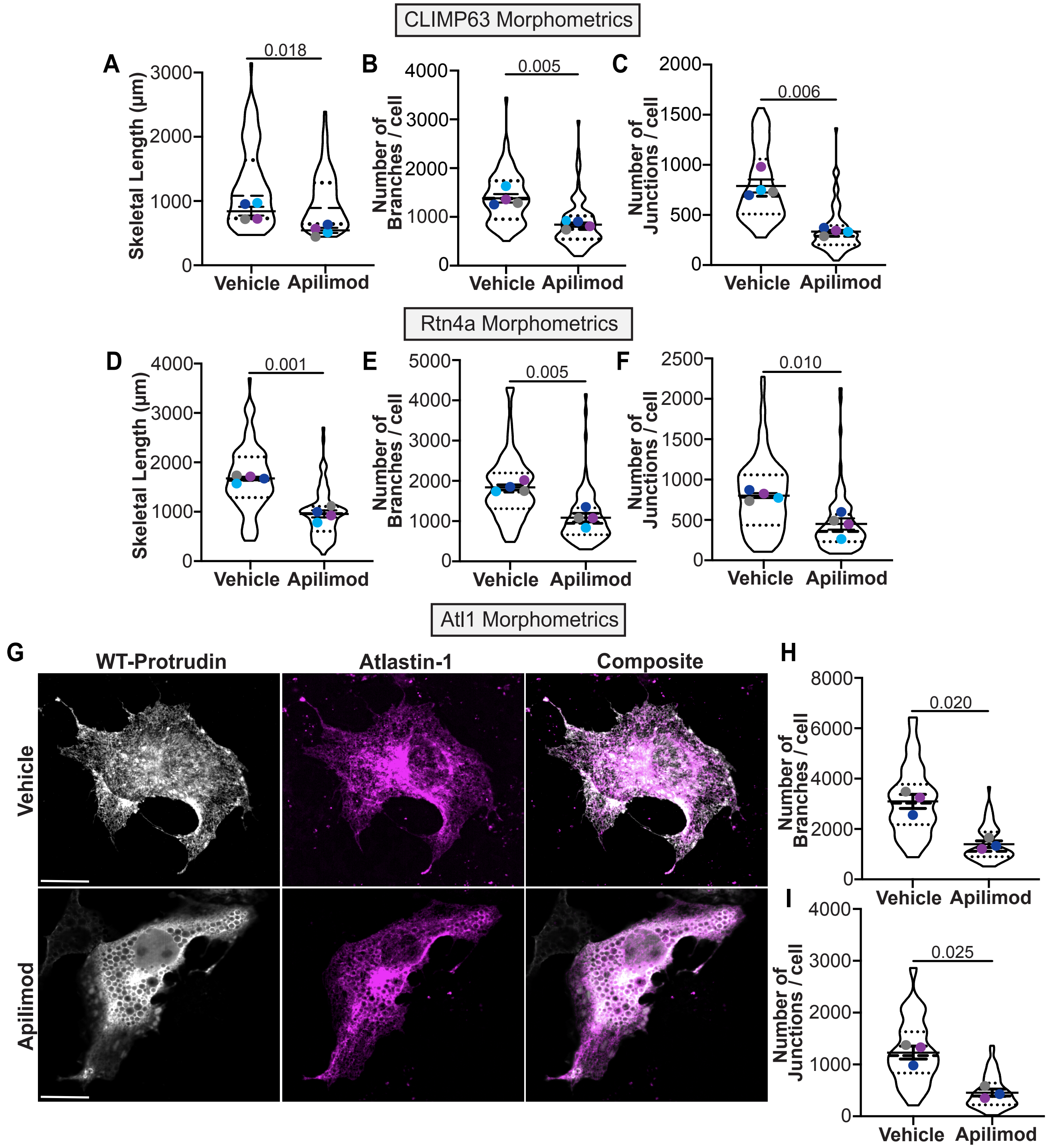
